# Genome-Wide Mapping of Human DNA Replication by Optical Replication Mapping Supports a Stochastic Model of Eukaryotic Replication

**DOI:** 10.1101/2020.08.24.263459

**Authors:** Weitao Wang, Kyle Klein, Karel Proesmans, Hongbo Yang, Claire Marchal, Xiaopeng Zhu, Tyler Borrman, Alex Hastie, Zhiping Weng, John Bechhoefer, Chun-Long Chen, David M. Gilbert, Nicholas Rhind

## Abstract

DNA replication is regulated by the location and timing of replication initiation. Therefore, much effort has been invested in identifying and analyzing the sites of human replication initiation. However, the heterogeneous nature of eukaryotic replication kinetics and the low efficiency of individual initiation site utilization in metazoans has made mapping the location and timing of replication initiation in human cells difficult. A potential solution to the problem of human replication mapping is single-molecule analysis. However, current approaches do not provide the throughput required for genome-wide experiments. To address this challenge, we have developed Optical Replication Mapping (ORM), a high-throughput single-molecule approach to map newly replicated DNA, and used it to map early initiation events in human cells. The single-molecule nature of our data, and a total of more than 2000-fold coverage of the human genome on 27 million fibers averaging ~300 kb in length, allow us to identify initiation sites and their firing probability with high confidence. In particular, for the first time, we are able to measure genome-wide the absolute efficiency of human replication initiation. We find that the distribution of human replication initiation is consistent with inefficient, stochastic initiation of heterogeneously distributed potential initiation complexes enriched in accessible chromatin. In particular, we find sites of human replication initiation are not confined to well-defined replication origins but are instead distributed across broad initiation zones consisting of many initiation sites. Furthermore, we find no correlation of initiation events between neighboring initiation zones. Although most early initiation events occur in early-replicating regions of the genome, a significant number occur in late-replicating regions. The fact that initiation sites in typically late-replicating regions have some probability of firing in early S phase suggests that the major difference between initiation events in early and late replicating regions is their intrinsic probability of firing, as opposed to a qualitative difference in their firing-time distributions. Moreover, modeling of replication kinetics demonstrates that measuring the efficiency of initiation-zone firing in early S phase suffices to predict the average firing time of such initiation zones throughout S phase, further suggesting that the differences between the firing times of early and late initiation zones are quantitative, rather than qualitative. These observations are consistent with stochastic models of initiation-timing regulation and suggest that stochastic regulation of replication kinetics is a fundamental feature of eukaryotic replication, conserved from yeast to humans.

## Introduction

The human genome is replicated from approximately 50,000 distinct initiation events in each cell cycle (Masai et al., 2010). However, identifying the location and firing timing of human replication origins is an ongoing challenge (Hyrien, 2015). A major part of this challenge is the inefficient and heterogeneous nature of mammalian origin firing (Berezney et al., 2000; Taylor, 1977). Even in cases where mammalian origins are thought to occur at relatively well-defined sites, they appear to fire with as little as 10% efficiency (Dijkwel et al., 2002); in the more common case where initiation appears to occur within broad initiation zones (IZs), spanning tens of kilobases (Hamlin et al., 2008), they fire with less than 1% efficiency (Demczuk et al., 2012). Therefore, population-based bulk approaches to mapping origins, which average origin-firing behavior across large populations of cells, must contend with low signal-to-noise ratios. As a result, although a number of bulk approaches have been used to map origins in the human genome (Mesner et al., 2013; Besnard et al., 2012; Cayrou et al., 2011; Foulk et al., 2015; Langley et al., 2016; Macheret and Halazonetis, 2018; Dellino et al., 2013; Petryk et al., 2016), there is little concordance among them (Langley et al., 2016; Mesner et al., 2013).

The heterogeneous nature of mammalian origins may be influenced by the non-site-specific nature of the mammalian origin recognition complex (ORC) DNA binding. Unlike budding yeast ORC, which binds a specific 17-bp sequence (Theis and Newlon, 1997; Eaton et al., 2010), mammalian ORC shows only a weak affinity for AT-rich sequences *in vitro* (Vashee et al., 2003; De Carli et al., 2018) and no discernible sequence preference *in vivo* (Miotto et al., 2016). G-quadruplex (G4) sequences have been proposed to function as origins (Cayrou et al., 2015; Valton et al., 2014; Prorok et al., 2019), but this hypothesis is controversial (Foulk et al., 2015; Miotto et al., 2016). A simple explanation for the observed patterns of ORC binding and replication initiation is that ORC binds non-specifically to accessible, nucleosome-free DNA (Lubelsky et al., 2011). In particular, ORC binding and replication initiation correlate well with DNase I accessibility across metazoan genomes (Miotto et al., 2016; Gindin et al., 2014b). Furthermore, MCMs can slide after loading, further dispersing sites of initiation (Gros et al., 2015; Edwards et al., 2002; Powell et al., 2015). For these reasons, we will refrain from referring to mammalian sites of replication initiation as origins but rather refer to specific instances of replication initiation as *initiation events* occurring at *initiation sites* and to areas enriched in initiation sites as *initiation zones*.

Although the specific locations of replication initiation are unclear, the general characteristics of mammalian sites of initiation have been described. Initiation is enriched in euchromatic promoter and enhancer regions, consistent with its correlation with accessible chromatin (Ganier et al., 2019; Petryk et al., 2016; Pourkarimi et al., 2016; Cayrou et al., 2015). Although initiation tends to be enriched in euchromatic regions, initiation must also occur in heterochromatin. Initiation in heterochromatin appears to be even more heterogeneous, making heterochromatic initiation sites even less well understood (Petryk et al., 2016; Cayrou et al., 2015).

Much effort has also been put into understanding the relative timing of DNA replication across the genome (Rhind and Gilbert, 2013; Fragkos et al., 2015; Marchal et al., 2019). The replication timing of a genomic region is primarily determined by the timing of initiation within that region. Based on population replication timing data, metazoan genomes can be divided into regions of similar local replication timing referred to as constant timing regions (Hiratani et al., 2008; Farkash-Amar et al., 2008; Hansen et al., 2010). Constant timing regions consist of smaller units whose replication timing can be regulated during development; these smaller units are called replication domains (Hiratani et al., 2008). Replication domains tend to align with topologically associated domains (TADs) and are thought to represent discrete functional units of the genome (Pope et al., 2014). The replication timing of replication domains correlates with the level of expression of the genes they contain and their chromatin modifications. However, whether the replication timing is a cause, a consequence, or an independent correlate of chromatin structure is a matter of debate (Rhind and Gilbert, 2013).

In budding yeast, where replication origin locations are well defined, origin firing is stochastic, with each origin firing with a specific probability, independent of neighboring origins (Czajkowsky et al., 2008; Yang et al., 2010; de Moura et al., 2010). Such stochastic firing leads to reproducible replication profiles at the population level, because more efficient origins are more likely to fire early and therefore, on average, have early replication times; by contrast, inefficient origins usually fire late or are passively replicated (Rhind et al., 2010). The heterogeneous and inefficient nature of metazoan replication initiation is also consistent with stochastic initiation. Simulations with a stochastic firing model, in which initiation is regulated only by a local-firing-probability function, faithfully reproduce experimental genome-wide replication timing profiles, suggesting that no deterministic timing program is required (Gindin et al., 2014b). Furthermore, if the local initiation rate is predicted by DNase I hypersensitivity, the simulation closely matches experimental results, consistent with the observed correlation between promoters and enhancers, which are DNase I hypersensitive, and initiation frequency (Gindin et al., 2014b). On the other hand, the reproducible replication timing of individual replication domains measured in single cells has led to the suggestion that replication within those domains initiates at defined times in most cells in the population (Dileep and Gilbert, 2018). Furthermore, neighboring initiation sites have been proposed to show both cooperative firing and lateral inhibition (Cayrou et al., 2011; Guilbaud et al., 2011; Löb et al., 2016), neither of which are consistent with strictly stochastic models. Therefore, whether metazoan initiation timing is stochastic or deterministic, or some combination of the two, is still very much an open question (Bechhoefer and Rhind, 2012).

A powerful solution to the problems of heterogeneity and low signal-to-noise ratios is single-molecule analysis, which allows the identification of sites of replication initiation on individual DNA fibers (Técher et al., 2013). Traditional single-molecule approaches—such as fiber autoradiography (Huberman and Riggs, 1968), DNA combing (Herrick and Bensimon, 1999; Anglana et al., 2003; Kaykov et al., 2016) and SMARD (Norio and Schildkraut, 2001) — have provided critical insight into the location and firing kinetics of mammalian replication origins. However, these techniques are restricted to the analysis of at most a few genomic loci. Nanopore-sequencing-based methods offer the potential to map replication genome-wide at high resolution (Müller et al., 2019; Georgieva et al., 2019; Hennion et al., 2020), but the current throughput of nanopore sequencing does not allow genome-wide analysis of metazoan genomes and current average read lengths are on the order of only 30 kb. Optical mapping technologies provide ample throughput on individual long (150 kb to 2 Mb) DNA fibers (Lam et al., 2012) and can be combined with nucleotide analog incorporation to map DNA replication (De Carli et al., 2018). However, the current approach relies on DNA replication in extracts (De Carli et al., 2018), precluding *in vivo* analysis of replication initiation.

We have developed Optical Replication Mapping (ORM), a single-molecule technique to investigate spatial and temporal distribution of origin firing that combines the Bionano Genomics approach to mapping long individual DNA molecules (Lam et al., 2012) with *in vivo* fluorescent nucleotide pulse-labeling (Wilson et al., 2016; Panning and Gilbert, 2005) to directly visualize sites of replication initiation within human cells. This approach affords us excellent signal-to-noise characteristics and deep, genomewide coverage, allowing us to identify initiation sites active in as few as 0.1% of human cells. We have used ORM to identify and analyze human sites of DNA replication initiation. We have further applied ORM to asynchronous human cells to obtain cell-type-specific replication profiles.

## Results

### Optical Replication Mapping of Early-Firing Human Initiation Sites

We mapped sites of early replication in the human genome by ORM, combining *in vivo* labeling of replication initiation with Bionano visualization and mapping of long individual DNA molecules (Lam et al., 2012, Figure 1A,B and Methods;). HeLa S3 cells were synchronized in mitosis by mitotic shake off and arrested at the beginning of S phase with aphidicolin. This protocol allows early initiation events but prevents elongation of replication forks from these initiation events and also prevents subsequent initiation events by activation of the intra-S-phase checkpoint (Feijoo et al., 2001). Cells were then electroporated with fluorescent ATTO647-dUTP, released from aphidicolin, allowed to recover overnight to complete replication, and harvested for analysis on the Bionano Saphyr platform. As a result, cells incorporate the electroporated fluorescent nucleotides in replication forks that initiated during the aphidicolin arrest and any that are initiated immediately after release. However, the transfected nucleotides are rapidly depleted, preventing incorporation at later initiation sites (Wilson et al., 2016). In our primary experiment, 87% of cells were synchronized in mitosis and, after release from aphidicolin, 94% incorporated the fluorescent label, with 100% of labeled cells showing an early-replication pattern of replication foci (Table S1, Figures S1A-C), indicating a high degree of synchrony and <1% of contaminating asynchronous S phase cells.

**Figure 1.**
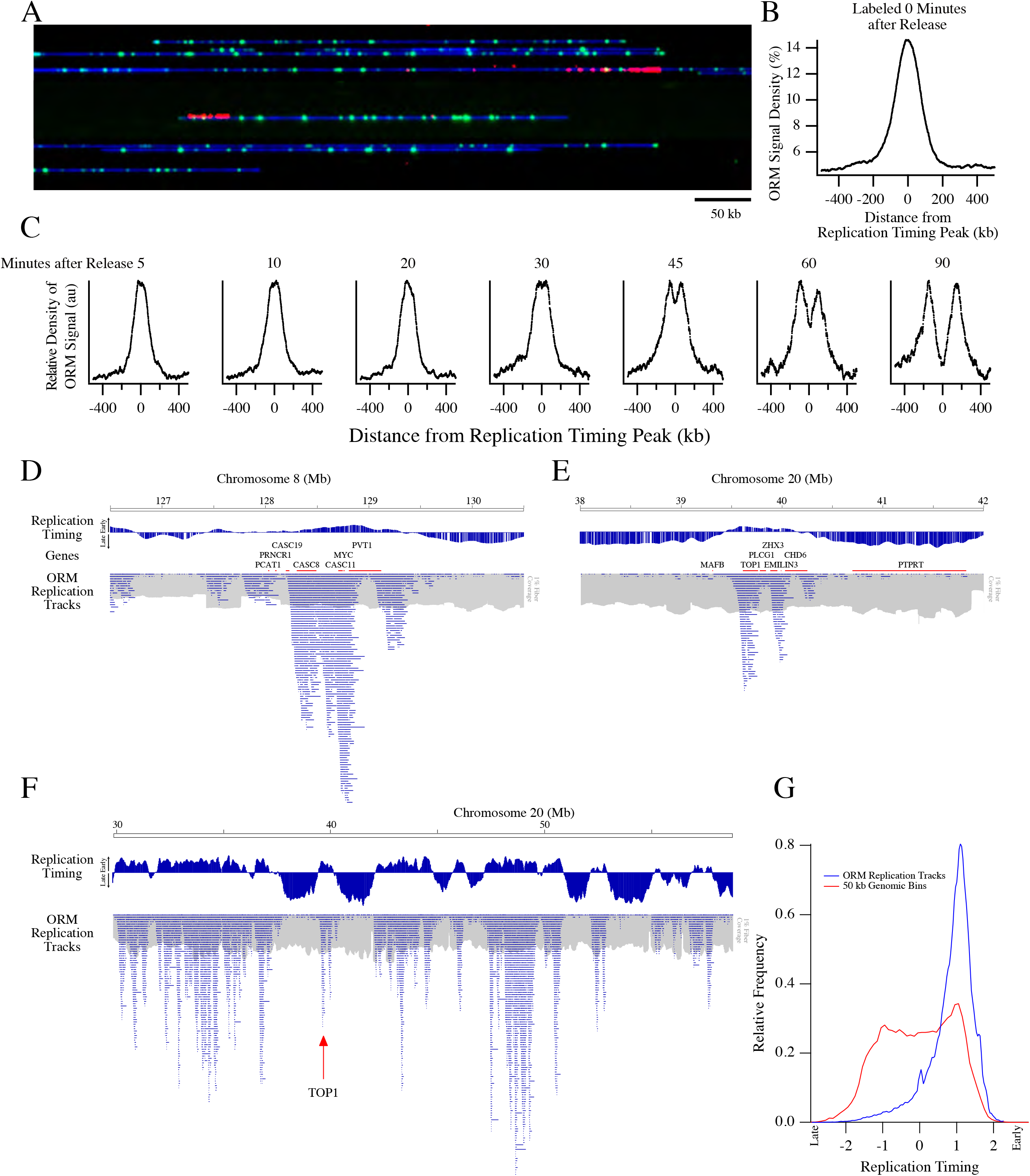
Optical Replication Mapping of Synchronized Cells Identifies Early Initiation Events. HeLa S3 cells were synchronized by mitotic shake off, arrested in early S-phase with aphidicolin, electroporated with fluorescent dUTP and released from the aphidicolin arrest, allowing fluorescent nucleotide incorporation around early replication initiation events. **A**) A representative image of the Bionano data with DNA in blue, Nt.BspQI restriction sites in green and incorporated nucleotides in red. The field of view is 750 kb. **B**) Enrichment of ORM signals from the combined 0-minute dataset around early replication timing peaks, those that replicate in the first quarter of S phase. **C)** Enrichment around early replication timing peaks of ORM signals from datasets in which cells were labeled between 5 and 90 minutes after release from synchronization. **D, E**) Distribution of replication tracks at two previously characterized human replication IZs: Myc (C) and Top1(D). The distribution of replication tracks in shown beneath the HeLa S3 replication timing profile (Hansen et al., 2010) and genes from the region. The distribution of total fibers in shown in grey at 1% the scale of the replication tracks. Discontinuities in the fiber density distribution, such as the one around megabase 127.5 in D, are probably caused by segmental duplication increasing the copy number of the region (see Figure S1E). Therefore, all analyses are normalized to fiber coverage depth. **F**) Distribution of replication tracks across the right arm of Chromosome 20, as in D. **G**) Distribution of replication timing of replication tracks compared to replication timing of each 50 kb bin in the genome.

We collected a dataset of 728 Gb across 2.9 million fibers ranging in length from 150 kb to 1.2 Mb, with an average fiber length of 250 kb and an average genome coverage of 206 fold (Table S1 and Figure S1D). On these fibers, we identified fluorescent incorporation signals and mapped them across the genome (Figure 1A, see Methods for details). To determine the technical reproducibility of our data, we collected it as three technical replicates. The replicates show high reproducibility, with the variation dominated by counting noise in infrequently labeled regions of the genome (r = 0.81, Figure S1F). To determine the biological reproducibility of our assay, we repeated the experiment three times and compared the signal distribution in the four biologically-independent experiments. The four replicates show high correlation (r = 0.83-0.86, Figure S1G), demonstrating low biological variation in our assay. Therefore, we combined the four experiments into one dataset of 11.8 million fibers ranging from 150 kb to 2.2 Mb in length, averaging 285 kb in length and constituting 1,109-fold coverage of the human genome (Table S1). The coverage is uniform, with a coefficient of variation of 12% (Figure S1E).

The replication-incorporation signal we detect is punctate in nature and uniform in intensity (Figures 1A and S2A). From these observations, we conclude that these incorporation signals consist of individual fluorescent nucleotides. Furthermore, the distributions of signal intensity (Figure S2A) and inter-signal distances (Figure S2B) are consistent with sparse labeling by a pool of fluorescent nucleotides that is depleted by replication, as expected for a one-time bolus of nucleotides delivered by electroporation. The distribution of the signals is predicted by inefficient, random incorporation of fluorescent nucleotides at an initial rate of about 1/900 thymidines that is depleted with a half length of about 75 kb (Figure S2B). Given the lengths of the replication tracks and the sparse nature of the signal within them, we estimate that we can localize initiation events with a resolution of about 15 kb (14.2±0.1 kb, see Methods for details).

The fluorescent signals in our initial ORM dataset are enriched in replication IZs, as identified in HeLa cells by replication timing peaks (Chen et al., 2011), consistent with our prediction that they represent nucleotide incorporation near sites of replication initiation (Figure 1B). To test whether the incorporation signals truly reflected initiation events, we modified our protocol to allow for a delay, ranging from 5’ to 90’, after aphidicolin release and before transfection (Table S1). We predicted that this approach would result in a gap between two symmetric populations of incorporation signals surrounding the replication initiation sites. As predicted, when we aggregate replication signals around early IZs, we see a time-dependent movement of incorporation tracks away from the IZs at an estimated rate of 1.65±0.31 kb per minute (Figure 1C), consistent with prior estimates of mammalian replication fork rates (Jackson and Pombo, 1998; Conti et al., 2007; Chagin et al., 2016). This result is consistent with the signal in our 0’ data representing initiation events and the signals after various release times representing forks moving away from such initiation events.

To facilitate visualization of labeled replication forks, we segmented the fluorescent-nucleotide incorporation signal into discrete replication tracks, identifying replication forks active during our labeling period (see Methods for segmentation details). In our combined 0’ dataset, we observed 977,746 replication tracks averaging 19.5±30.9 kb in length. The observed average track length of 20 kb is shorter than the 75 kb that we estimate is replicated during the labeling period (Figure S2B) due to the sparsity of the labeling. In addition, because of the sparse labeling, 58% of our replication tracks are labeled with only one signal, further reducing the average observed track length. To test whether these solo signals are sparsely labeled replication tracks or spurious background noise, we calculated the enrichment of solo signals at early replication sites. Solo signals are enriched at such sites, demonstrating that they, too, mark regions of replication initiation (Figure S2C), and that noise, if any, is very low with this method. Although many of the tracks are labeled with only one signal, few are not labeled at all. Our modeling of the rate of label incorporation and label depletion suggest that over 90% of initiation events in early S-phase will incorporate at least one label within 15 kb of replication (Figures S2A and B and Supplemental Mathematical Methods Section 4). Therefore, almost every early-initiation event will be visualized at a resolution of 15 kb.

For fibers on which we identify more than one track, the average distance between track centers is 111±78 kb (Figure S2D), consistent with previous measurements (Cayrou et al., 2011; Jackson and Pombo, 1998). We find enrichment of replication tracks around previously identified sites of human replication initiation (Figure 1D,E). In particular, we examined the distribution of replication tracks around the MYC and Top1 loci, both characterized as early-firing origins in the human genome (Tao et al., 2000; Keller et al., 2002), and found a pronounced enrichment at these loci, particularly in the intergenic regions surrounding these genes. More generally, replication tracks are enriched in the earliest replicating parts of the genome, as defined by replication timing profiling (Hansen et al., 2010), as expected for early-initiation events (Figures 1F,G). Interactive display of the ORM data is available on the ORM Browser <http://orm.nucleome.org> (Figure S3, Zhu et al. in prep, personal communication, J. Ma).

We observe 0.97 million initiation events in our combined 0’ dataset. From the median incorporation track length of 75 kb (estimated from our modeling of label-incorporation kinetics, which compensates for the sparseness of labeling, Figure S2B), we infer that 73 Gb of DNA is labeled, which is ~2% of the 3.36 Tb in the dataset. Therefore, we are labeling the DNA replicated in the first ~2% of S phase and during that time, about 1000 initiation events occur per genome (977,746 initiation events observed in 1,109 genome equivalents).

### Determination of Genome-Wide Replication Kinetics in Asynchronous Cells

The experiments described above provide unprecedented single-molecule mapping of initiation events at the onset of S phase. However, most cell types are not amenable to such precise cell-cycle synchronization. To map replication kinetics in unperturbed cells, we labeled asynchronous HeLa cells and H9 human embryonic stem cells, and mapped replication incorporation tracks by ORM (Figure 2A). We identified 412,113 replication tracks in two biologically-independent HeLa replicates totaling 1.4 Tb of data and 299,595 tracks in one H9 dataset totaling 738 Gb of data (Table S1). Using the same analysis as for our synchronous datasets, we infer a similar nucleotide-labeling frequency of 1/1025 and 1/850 thymidines, respectively. The replication tracks average 23.9±35.5 kb in length in the HeLa data and 27.5±40.4 kb in length in the H9 data, which is comparable to the length of tracks in the synchronized data. Thus, forks released from aphidicolin arrest synthesize at about the same rate as untreated replication forks. The tracks are uniformly distributed across the genome, as predicted for asynchronous replication forks, with an average density of 1.3±0.5%. In particular, in contrast to our synchronous dataset, and as expected, we see no enrichment at replication timing peaks in early- or late-replicating regions (Figure 2B).

**Figure 2.**
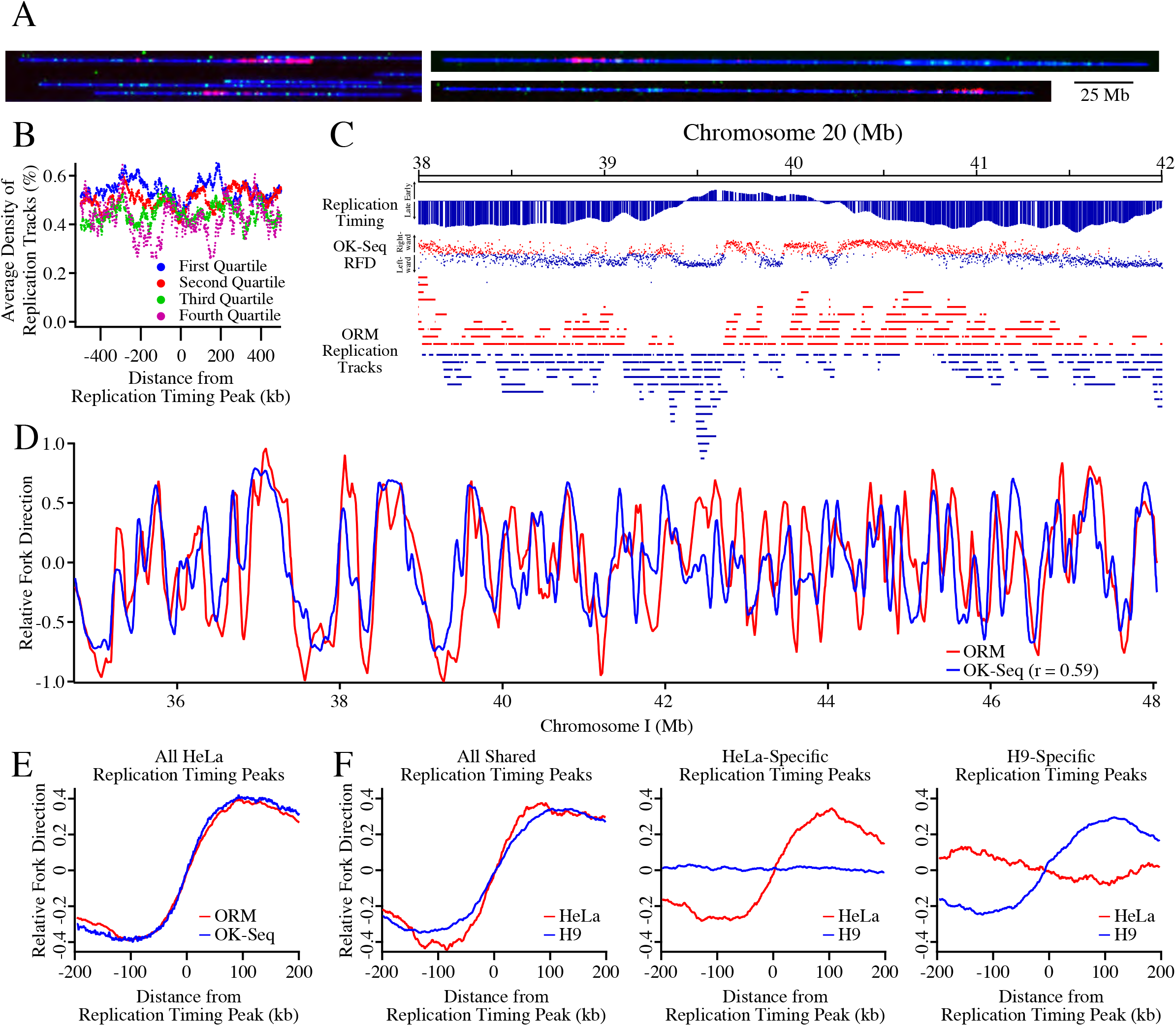
Optical Replication Mapping of Asynchronous Cells Identifies Cell-Type-Specific Genome-Wide Replication Kinetics. A) Asynchronous HeLa S3 and H9 human embryonic stem cells were electroporated with fluorescent dUTP, allowing fluorescent nucleotide incorporation into ongoing replication fork. **B)** The density of ORM signals from the HeLa asynchronous dataset around replication timing peaks separated into replication timing quartiles, defined by average replication timing (S50, see Methods). **C**) Labeled ORM replication forks, with leftward-moving forks colored blue and rightward-moving forks colored red, are depicted below the replication timing profile of the Top1 region and its OK-seq relative fork direction (RFD) data, with blue dots indicating leftward replication, red dots rightward, and the vertical position of the dot, between −1 and 1, indicating the fraction of replication moving in that direction. The polarity of ORM replication forks was determined by calculating their fork direction index (FDI, see Methods). **D**) The RFD calculated from ORM and OK-seq data across 7 Mb of Chromosome 1. E) The enrichment of HeLa fork polarity, as measured by RFD, around early replication timing peaks. **F**) The enrichment of HeLa and H9 fork polarity around cell-type specific and cell-type general early replication timing peaks.

In order to use this data to infer replication kinetics, it is necessary to determine the polarity of each incorporation track, so that the direction of each mapped replication fork is known. To determine track polarity, we took advantage of the fact that, as the labeled nucleotides are consumed, the label density diminishes, a strategy that has been used in other fiber-analysis approaches (Müller et al., 2019; Huberman and Riggs, 1968; Hennion et al., 2020). We calculated a fork direction index (FDI) as the ratio of integrated fluorescent signal in the left half of the track to that in the right half, for all tracks with 3 or more signals.

As expected, ORM replication tracks tend to be oriented in the direction of replication inferred from the slope of HeLa cell replication timing profiles and from HeLa cell Okazaki fragment mapping (Hansen et al., 2010; Petryk et al., 2016) (Figures 2C,D). Furthermore, genome-wide analysis of replication fork polarity around replication timing peaks shows that the polarity signal in ORM data is comparable to that in published OK-seq data (r = 0.59, Figures 2D,E). Importantly, the polarity signal in ORM data is celltype specific (Figure 2F). These results show that ORM data can be used to characterize replication kinetics in unsynchronized cells, demonstrating its applicability to any cells that can be pulse labeled with fluorescent nucleotides.

### Genome-Wide Mapping of Early-Firing Human Initiation Zones

To map the genome-wide distribution of early-firing human initiation sites, we analyzed the incorporation signal in our combined 0’ dataset and identified 4,930 IZs, defined as peaks of replication signal density (Figure 3A, see Methods for details). The IZs are mostly between 20 and 40 kb, with an average length of 32.3±15.9 kb (Figure S2E), comparable to IZs mapped by OK-seq (Petryk et al., 2016). The initiations zones cover about 5.3% of the genome and contain about 19.8% of the ORM signal. Much of the rest of the ORM signal is directly adjacent to defined IZs, but much is distributed elsewhere at lower density, particularly in late-replicating regions on the genome (Figure 3A). Plotting the initiation efficiency of 50 kb genomic bins show that there is a continuous distribution of initiation efficiency between the higher-efficiency regions we define as IZs and lower-efficiency regions, with no obvious break between them (Figure S2F).

**Figure 3.**
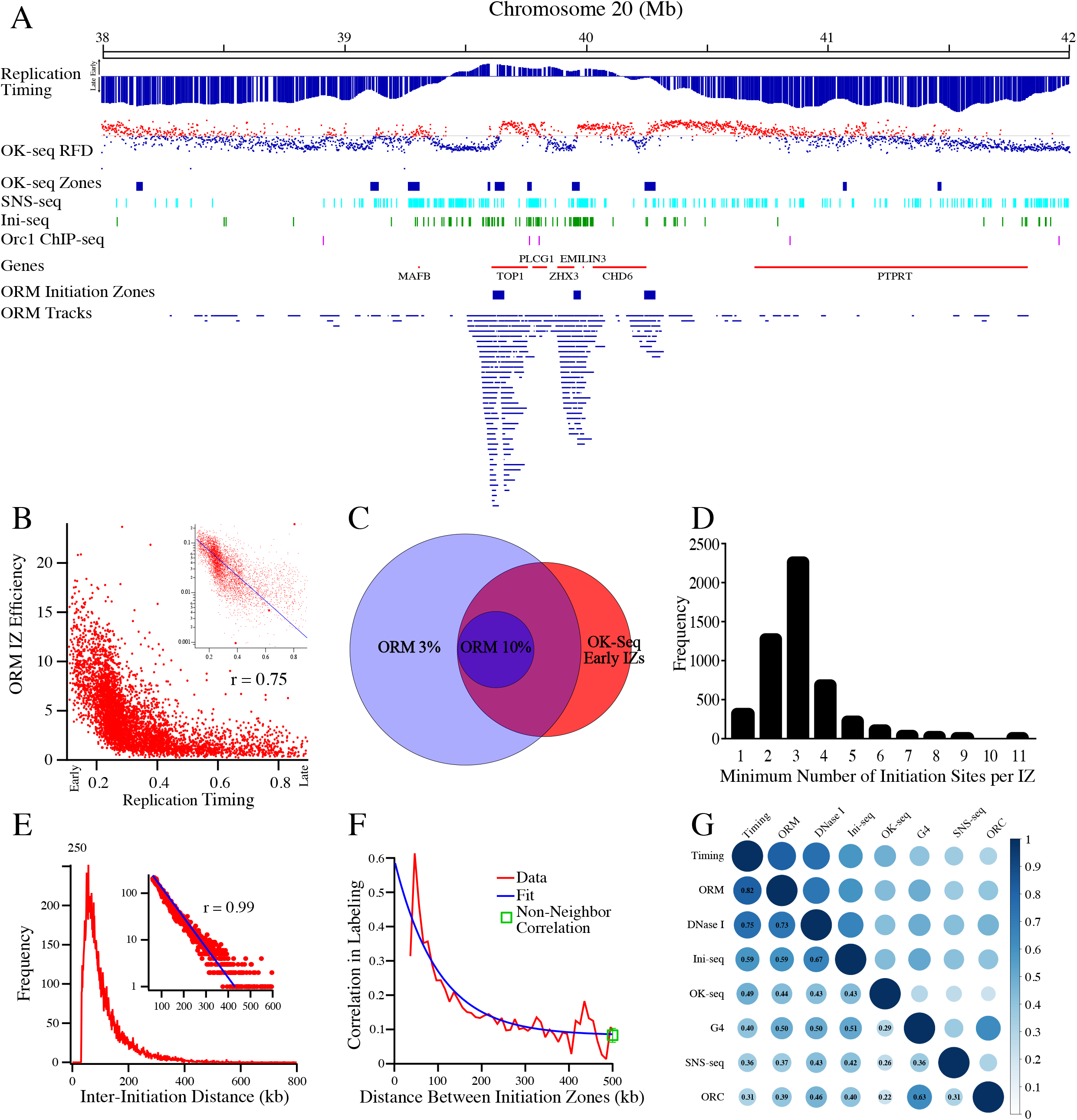
Mapping of Human Replication Initiation Zones. **A**) A comparison of ORM tracks, ORM IZs, OK-seq RFD (as in Figure 2C), OK-seq IZs (Petryk et al., 2016), SNS-seq (Picard et al., 2014), Ini-seq (Langley et al., 2016) and Orc1 ChIP-seq (Dellino et al., 2013) at the Top1 locus. **B**) The correlation between ORM initiation efficiency (fraction of labeled fibers/IZ) in the combined 0-minute dataset and replication timing (S50) in the 4,930 ORM IZs. Spearman rank correlation coefficient between the two datasets is 0.75. Inset is the same data on a semi-log plot fit with an exponential curve. **C**) A Venn diagram of the overlap between OK-seq early IZs and 50 kb windows in the genome that have ORM initiation efficiencies of at least 3% or 10%. **D**) The estimated minimal number of initiation sites per IZ. The estimate was made by calculating minimum number of replication tracks in each IZ whose centers are more than 15 kb apart. Most of the IZs for which all track centers are within 15 kb of each other are small and contain few replication tracks (Figure S4B). **E**) The distribution of the inter-initiation distances in the combined 0’ dataset measured as the distance from the middle of neighboring replications tracks for fibers that have multiple tracks. The average of the distribution is 111 kb and the mode of distribution is 57 kb. Inset: The distribution, plotted on a log y-axis from 60 to 600 kb, fit to an exponential curve (r = 0.99). The exponential distribution of the interreplication-track distances indicates that the distribution of initiation events on this length scale is random. **F**) The correlation between the probability of replication at neighboring IZs. The non-neighbor correlation (0.06±0.01, mean±s.e.m.) is the average correlation between any two IZs on one fiber more than 200 kb apart, irrespective of the number of intervening IZs. **G**) A matrix depicting the correlation between initiation sites predicted by ORM, replication timing (Chen et al., 2010), OK-seq (Petryk et al., 2016), SNS-seq (Picard et al., 2014), Ini-seq (Langley et al., 2016) Orc1 ChIP-seq (Dellino et al., 2013), DNase I hypersensitivity (Bernstein et al., 2012) and G4 motifs (Puig Lombardi et al., 2019).

The median genome-wide inter-initiation-zone distance is 273 kb. However, since our dataset is biased towards early replicating IZs, we suspect that we underestimate the number of IZs in late-replicating parts of the genome. Consistent with that contention, limiting our analysis to the earliest quarter of the genome leads to a median inter-initiation-zone distance of 168 kb. The inter-initiation-zone distances in the other three quarters of the genome are 257, 258 and 337 kb, respectively. We suspect that the lower density of observed IZs in the later-replicating regions of the genome results from having less data rather than fewer initiation events in these regions (Figure 1G).

A major advantage of ORM is that it records the number of fibers that both contain and do not contain replication tracks, allowing for the calculation of the absolute efficiency with which replication initiates at each locus in the genome. Although such an analysis is possible with prior methods, it is considerably more difficult and thus rarely reported (Demczuk et al., 2012). Such an analysis shows that early initiation is quite heterogeneous. Examining the IZs with average replication times in the first quarter of S phase (S50 < 25%), we find that they initiate replication, on average, on only 7.5% of molecules (Figure 3B). Even the 5% earliest IZs are active on only 11% of molecules. We therefore conclude that, in aphidicolin arrested cells, replication initiates in only a stochastically-selected subset of IZs, including only 11% of the most efficient early IZs.

We investigated the frequency of early initiation of replication across the genome, examining all 4,930 ORM IZs. We find that the local efficiency of initiation in these zones is generally below 15%, with occasional efficiencies up to 20% and a median efficiency of 3.7% (Figure 3B). All of the IZs with efficiencies of more than 10%, and 99% of the zones with efficiencies of more than the median of 3.7%, are in early-replicating regions of the genome (Figure 3B), consistent with the results in Figure 1G. We also find a strong concordance between the IZs identified by ORM and those identified by OK-seq (Petryk et al., 2016). In particular, 68% of the 6,166 early IZs identified by OK-seq show at least 3% initiation efficiency by ORM and 100% of loci that show greater than 10% efficiency by ORM are identified by OK-seq as IZs (Figure 3C).

Visual inspection of the distribution of replication tracks suggests that initiation in human cells is not concentrated at discrete, frequently-used origins. Instead, initiation appears to be distributed across broad IZs, as has been observed at specific loci and in other genome-wide analyses (Petryk et al., 2016; Mesner et al., 2013; Besnard et al., 2012; Hamlin et al., 2008; Dijkwel et al., 2002; Demczuk et al., 2012; Lubelsky et al., 2011; Tao et al., 2000). Even the IZ at the Top1 locus appears to have initiation events distributed over tens of kilobases (Figures 1E, S4A). However, given the sparse labeling of our replication tracks, we cannot identify initiation sites with resolution higher than about 15 kb. Nonetheless, 93.5% of IZs contain at least two incorporation tracks with centers that are more than 15 kb apart, suggesting that these IZs contain at least two, and possibly many more, initiation sites (Figures 3D and S4A,B). Further, the 6.5% of initiation zones that do not have obviously non-overlapping replication initiation sites tend to be smaller (16 kb v. 33 kb), less efficient (1.2% v. 4.5%) and later replicating (S50 0.55 v. 0.32) than those that do (Figure S4B). Furthermore, a reanalysis of 66 potentially discrete initiation site identified by OK-seq is consistent with the conclusion that there are no isolated, discrete initiation sites in the human genome (Figure S5).

Another way to distinguish between initiations that occur at a single site and those that are distributed across a broad region is to examine the distribution of ORM signals across the IZs, the expectation being that a single initiation site will have narrower distribution than distributed initiation sites. We measured the signal at the edges of IZs as a fraction of the signal at the middle of the IZ and fit to that data two models, one based on a single initiation site and another based on a uniform distribution of initiation sites across the IZ (Figure S4C). For IZs less that 55 kb long, which includes over 90% of IZs, the distribution signal is consistent with a uniform distribution of initiation sites throughout the IZs. Taking all of our data together, we find no evidence of isolated or high-efficiency initiation sites in the human genome.

### Initiation Events in Neighboring Initiation Zones are not Clustered

Regardless of how initiation events are locally distributed, it is important to know whether local initiation events are coordinated. A popular hypothesis in the field is that initiation events are clustered on chromosomes, such that initiation events will occur simultaneously in neighboring IZs (Fragkos et al., 2015). However, this hypothesis has never been rigorously tested, because the single-molecule experiments that have identified multiple initiation events on individual chromosomes have never quantitated their frequency relative to single initiation events in same IZ (Jackson and Pombo, 1998; Cayrou et al., 2011; Lebofsky et al., 2006; Huberman and Tsai, 1973; Blow et al., 2001; Marheineke and Hyrien, 2004), which is necessary to determine if multiple initiation events happen more often than would be expected by chance.

With our deep, single-molecule data on long (up to megabase-length) fibers, we can quantitatively test whether initiation events in neighboring IZs are correlated. A simple, unbiased test for correlation between events is to determine whether the spacing between the event deviates from an exponential distribution. Items distributed randomly on a line have an exponential distribution of inter-item distances (Birnbaum, 1954); therefore, correlated clusters of initiation events would deviate from an exponential distribution of inter-initiation distances. We measured 18,275 inter-initiation distances on 13,769 fibers containing multiple initiation events (Figure 3E). We measure few initiation events less than 60 kb apart, due to the facts that the tracks themselves average 40 kb in length and that early IZs have a median distance of 168 kb. However, above 60 kb, there is an exponential distribution of inter-initiation distances (r = 0.99, Figure 3E, inset). This distribution indicates that, on this length scale, initiation events are distributed randomly.

As an orthogonal test of whether initiation events in neighboring IZs are positively or negatively correlated, we compared the frequency of labeling at each IZ with the frequency with which its neighboring IZs were labeled on the same fiber (Figure 3F). Neighboring IZs can be co-labeled as a result of individual initiation in each zone or of initiation in one zone leading to passive incorporation of label in a neighboring zone by the same replication fork. Correlation can also arise because of heterogeneous uptake of labeled nucleotides. Basically, if one cell has a very high level of nucleotide uptake, all IZs (and any two neighboring IZs, in particular) will be more likely to incorporate label, making them appear to be correlated (see Methods for a more detailed explanation). Therefore, we fit a model to our initiation-zone co-labeling data that takes into account the previously determined incorporation track length (Figure S2B) and nucleotide uptake heterogeneity (Figure S1C), and assumes that individual initiation events are independent. This model fits the data well, suggesting there is no measurable positive or negative correlation between initiation in neighboring zones (Figure 3F). Furthermore, any apparent correlation due to heterogeneity of nucleotide uptake would be seen among all IZs in a cell, not just neighboring ones. Thus, we also calculated the correlation coefficient among all IZs on each fiber, not just the neighboring ones, and found a coefficient similar to that of neighboring IZs separated by more than 200 kb, consistent with the interpretation that the low level of residual correlation seen at longer distances is a global effect, not a local spatial correlation (Figure 3F). Finally, it is worth noting that the level of correlation that we ascribe to heterogeneous label uptake is only about 10%, much lower than proposed in synchronous-firing models.

### Comparison of Different Origin Mapping Datasets

We compared our optical replication mapping data to four published genome-wide HeLa replication-initiation-mapping datasets: OK-seq (Petryk et al., 2016), SNS-seq (Picard et al., 2014), Ini-seq (Langley et al., 2016) and Orc1 ChIP-seq (Dellino et al., 2013). By visual inspection, replication tracks appear to be enriched around the IZs identified by OK-seq (Figure 3A). Across the genome, we see colocalization of OK-seq IZs and ORM signal. We see similar colocalization of ORM signal with Ini-seq signal, a cell-free initiation mapping approach and, to a lesser extent, Orc1 ChIP-seq peaks and SNS-seq peaks, which maps initiation events by sequencing short nascent strands produced by replication initiation (Figure 3A). To determine whether the apparent co-localization of initiation mapping data is robust, and to quantify its extent, we measured the correlation between the five data sets (Figure 3G). We find that ORM replication tracks correlate well Ini-seq (r = 0.59), to a lesser extent with OK-seq (r = 0.49), and even less well with SNS-seq (r = 0.36) and Orc1 ChIP-seq (r = 0.31). These correlations are further confirmed by ROC analysis (Figure S6A).

### Characterization of Initiation Zones Mapped by ORM

To characterize the genomic character and chromatin context of our IZs, we investigated the enrichment of various genomic and epigenomic annotations, all derived from HeLa cells (Figure 4). Comparing our IZs with various chromatin states from the ENCODE project (Bernstein et al., 2012; Ernst and Kellis, 2012), we find that they are enriched in enhancers and low activity intergenic regions and depleted in transcription units and polycomb-repressed chromatin (Figure S7A). Examining specific histone modifications, our IZs are enriched in H2AZ, which has been implicated in ORC binding, and enhancer-specific modifications, such as H3K4me1 and H3K27ac, and they are depleted in, although adjacent to, transcription-elongation-associated modifications, such as H3K79me2 and H3K36me2 (Figures 4 and S7B,D), in agreement with previous studies (Petryk et al., 2016; Long et al., 2020; Pourkarimi et al., 2016). Consistent with the enrichment of initiation events in enhancer regions and their depletion in transcribed regions, IZs are enriched in DNase I hypersensitive sites and depleted in RNA pol II (Figures 4 and S7B,D).

**Figure 4.**
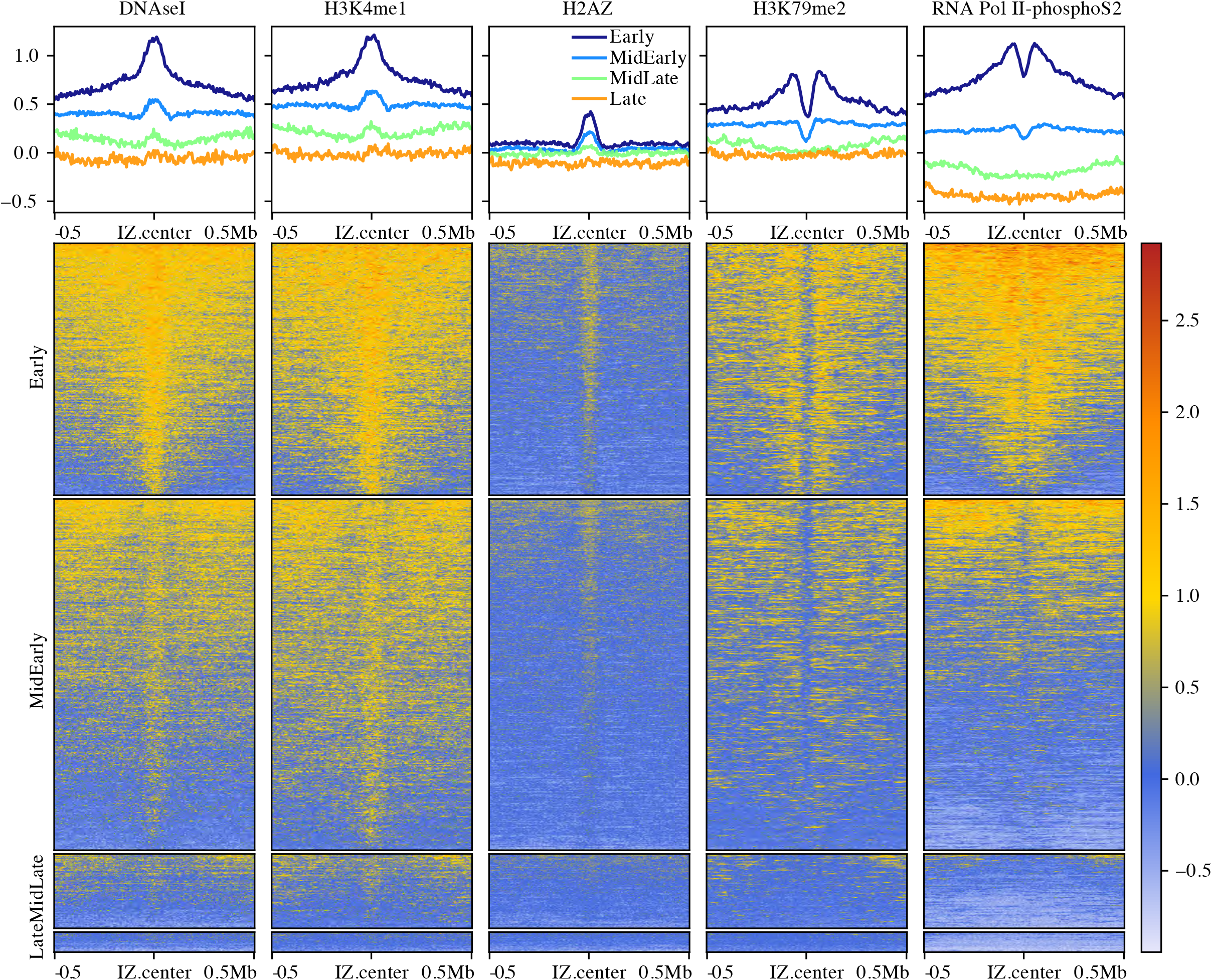
Characterization of Human Initiation Zones. Enrichment of histone modifications and other genomic features relative to ORM IZs, separated into replication timing quartiles. The upper panels show the genome-normalized relative signal around all IZs. The lower panels show heat maps of the same signal at each IZ. The y-axis scale for the upper panels and the heat-map scale for the lower panels are the same genome-normalized enrichment in relative units.

Although metazoan ORC has only a weak affinity for AT-rich sequences *in vitro* (Vashee et al., 2003; De Carli et al., 2018) and no discernible sequence preference *in vivo* (Miotto et al., 2016), sequences that form G4 quadruplex structures have been proposed to correlate with initiation sites in vertebrate cells (Cayrou et al., 2015; Langley et al., 2016; Valton et al., 2014; Prorok et al., 2019). We tested whether the IZs we identified are enriched in G4 motifs, using a GC-content-adjusted background model to avoid detecting G4 enrichment as a trivial consequence of the GC-rich nature of enhancers. Of the 737,735 G4 sequences computationally identified in the human genome (Puig Lombardi et al., 2019), 68,585 are found in IZs, which is not more than would be expected by chance, given their GC-rich nature (partial correlation r = −0.0513). Moreover, GC-rich regions (with similar GC% as IZs) outside of IZ are no less likely to contain G4 motifs (131,114 out of 225,710, 58.1%) than GC-rich regions within IZs (10,889 out of 18625, 58.4%, Fisher’s Exact Test p = 0.61). Therefore, although we find that IZs are GC rich (Figure S7C), as previously reported (Xu et al., 2012; Cayrou et al., 2015), they appear to contain G4 motifs as a consequence of their GC richness, but do not appear to be enriched for G4 motif DNA sequences, per se.

### Early Initiation Events in Late-Replicating Domains

Although 91% of our early initiation events are in early-replicating regions of the genome, 9% occur in late-replicating domains (Figures 1F, 3B). By visual inspection, the replication tracks in late-replicating parts of the genome appear to be in the locally earliest-replicating parts of the late-replicating regions (for instance at 56 and 58 Mb in Figure 1F), suggesting that they reflect the location of the IZs normally used to replicate these regions and not contaminating asynchronous cells. To test this suggestion, we calculated the enrichment of replication tracks around IZs identified as replication timing peaks (Chen et al., 2011) grouped into four quartiles, defined by average replication timing (S50). We find significant enrichment of replication tracks IZs in first three quartiles, (but no enrichment in the fourth quartile, which contains only 3% of the genome-wide replication timing peaks), demonstrating that our ORM signal in late-replicating regions is associated with IZs (Figure 5A). Moreover, the signal-to-noise ratios in the first three quartiles is similar (SNR = 3.2-2.8, Figure 5A), demonstrating that, although there is much less signal in the third quartile, it is enriched over background to the same extent as in the earlier quartiles. In contrast, no enrichment is seen in any quartile in asynchronous cells (Figure 2A). These results confirm that the enrichment of early initiation events we see at late-replicating IZs cannot be attributed to contaminating asynchronous background signals and, therefore, presumably results from bona fide early initiation events in IZs that, on average, replicate late in S phase.

**Figure 5.**
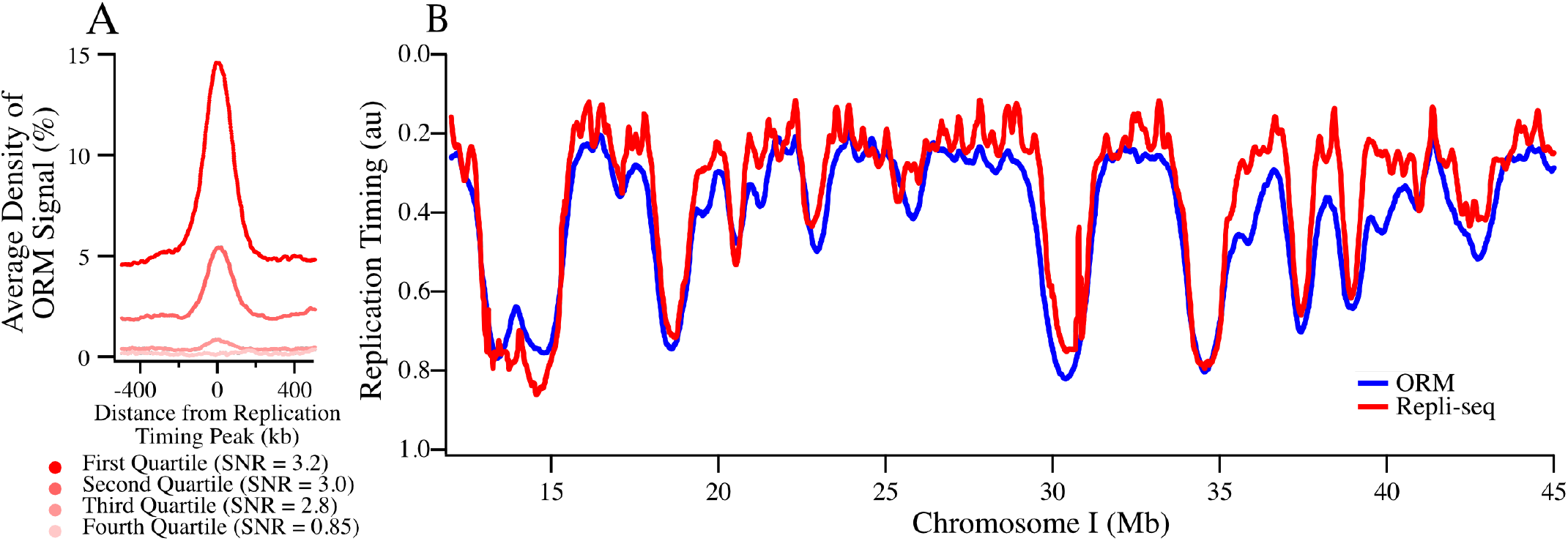
Probability of Early Initiation Predicts Replication Kinetics Genome-Wide. **A**) Enrichment of ORM signals from the combined 0-minute dataset around replication timing peaks separated into replication timing quartiles. Signal-to-noise ratio (SNR) was calculated as maximum signal over average signal between 400 and 500 kb away. **B**) Comparison between experimentally determined replication timing (S50) and replication timing predicted from ORM data using a stochastic model (r = 0.85, Gindin et al., 2014a).

### Computational Modeling Suggests that Early- and Late-Replicating Initiation Zones are Distinguished Only by Relative Initiation Probability

The distribution of early initiation events we observe—frequent firing in early-replicating IZs and rare firing in late-replicating IZs—is reminiscent of the stochastic regulation of origin-firing timing observed in budding yeast (Czajkowsky et al., 2008; Yang et al., 2010; de Moura et al., 2010). Models of stochastic replication timing posit that the only difference between early- and late-firing origins is their relative probability of firing, and that this relative probability does not change during S phase. More-efficient origins are more likely to fire first and thus, on average, have earlier firing times, but sometimes they still fire late, whereas less-efficient origins are less likely to fire early and thus, on average, have later firing times, but do sometimes fire early (Rhind et al., 2010).

Such models predict that, because the relative probability of origin firing does not change during S phase, measuring the relative probability of initiation in early-S phase is sufficient to determine the kinetics of initiation for all IZs throughout S phase. If this prediction holds true, we should be able to calculate the complete replication timing profiles of cells, even though we have assayed firing probabilities in only the first 2% of S phase. To test this prediction, we used Replicon, a stochastic replication simulator that calculates replication profiles from a so-called Initiation Probability Landscape (IPLS), the probability of initiation at each point in the genome (Gindin et al., 2014a). We find that we can, indeed, accurately recapitulate genome-wide replication timing profiles with our ORM data (r = 0.85, Figure 5B). Further, our ORM data predicts replication timing better than other initiation data sets, such as, OK-Seq (r = 0.81), Ini-seq (r = 0.69) or SNS-seq (r = 0.48, Figure S6B). ORM predicts replication kinetics equally as well as H3K4me1 (r = 0.85), which marks open enhancer and promoter chromatin, and DNase I hypersensitivity (r = 0.85), which had previously been identified as the best predictor of initiation (Gindin et al., 2014a). These result support the hypothesis that the primary difference between early- and late-replicating IZs is their relative initiation probability, influenced by chromatin accessibility, and not a qualitative difference in their initiation times.

## Discussion

We have developed Optical Replication Mapping (ORM), which, for the first time, allows genome-wide, high-throughput, single-molecule replication mapping in human cells. The approach is based on *in vivo* labeling of replication forks and *in vitro* detection and genomic mapping of incorporated label on long, single DNA fibers using the Bionano Saphyr platform. Single flow cells using the current mapping approach yielded an average 150-fold coverage of the human genome on fibers from 150 kb to 2 Mb, with an average length of 300 kb (Table S1). Altogether, we have collected 2,550-fold coverage with an average fiber length of 284 kb. Our method is similar to the recently described HOMARD approach, which was used to map DNA replication in *Xenopus* egg extracts (De Carli et al., 2018).

### Direct, single molecule, measurements of initiation frequency

Detection of DNA synthesis on individual DNA fibers is the most direct way to localize sites of DNA replication. Since the preponderance of the literature describes low-frequency, heterogeneous selection of replication initiation sites in mammalian cells, single-molecule measurements may be the only means to accurately map the relative frequencies of initiation events in the human genome. However, to date, single-molecule replication-mapping technologies, such as DNA combing and SMARD, which have throughput of a few hundred fibers, have provided insufficient depth of coverage for metazoan genomewide analyses. Recently, nanopore sequencing has been used to map replication incorporation tracks genome-wide on single DNA fibers, but the current technology only allows for relatively small datasets of relatively short fibers (Georgieva et al., 2019; Müller et al., 2019; Hennion et al., 2020). The largest dataset from a recent study describing the DNAscent approach to nanopore-based replication mapping in budding yeast was 3.8 Gb (the equivalent of just over 1x coverage of the human genome) with an average fiber length of 32 kb (Müller et al., 2019). Therefore, ORM, for which current typical datasets are 150-times larger, provides a unique opportunity for single-molecule, genome-wide mapping of metazoan replication at high coverage on long fibers.

Using ORM, we have mapped the replication initiation events occurring in the first 2% of S phase in aphidicolin-synchronized HeLa cells. These initiation events, as others have reported (Petryk et al., 2016; Long et al., 2020; Pourkarimi et al., 2016; Ganier et al., 2019; Cayrou et al., 2015), are enriched in the enhancer regions of active genes and are enriched in active chromatin modifications, in particular DNase I HS, H2AZ and H3K4me1 (Figure 4). However, unlike previous reports, we can directly estimate the frequency with which initiation occurs in each IZ because we measure all fibers, both labeled and unlabeled. We find that the distribution of early initiation events is heterogeneous. Although 99% of early IZs predicted by OK-seq analysis are detected in our ORM data, only 6% of them, on average, initiate at the beginning of S phase in any given cell. Furthermore, although most of the initiation events that we observe are in early-replicating IZs, a small but significant minority are in late-replicating IZs, and the frequency of initiation is correlated with the replication timing of the IZ (Figure 3B). This observation is not due to contamination of our synchronized G1/S cells with non-synchronized cells because the signal in asynchronous cells shows no enrichment at specific sites, let alone previously mapped late initiation sites (Figure 2A). These results suggest that, in any given cell, a small number of potential initiation sites are stochastically selected from a large number of potential initiation sites, and that the probability of a site being selected is correlated with the replication time of that site, with early-replicating sites having a higher probability of initiating than late-replicating sites (Figure 3B).

We have mapped replication initiation sites across the genome with over 1,000-fold genome coverage, allowing us to identify initiation sites used in as few as 0.1% of cells. The initiation events that we map appear to be spread over broad IZs, consistent with previous observations (Petryk et al., 2016; Mesner et al., 2013; Besnard et al., 2012; Hamlin et al., 2008; Dijkwel et al., 2002; Demczuk et al., 2012; Lubelsky et al., 2011; Tao et al., 2000; Anglana et al., 2003). However, the sparse labeling and relatively long length (~40 kb) of our incorporation tracks allow us estimate the location of initiation sites at a resolution of only about 15 kb. Nonetheless, in 93.5% of IZs, we observe incorporation track centers that are more than 15 kb apart, suggesting that they initiated from distinct sites (Figures 3D and S4). Furthermore, the distribution of signal across IZs is not consistent with a single initiation site per zone (Figure S4C). We cannot distinguish between frequent, discrete, origins scattered throughout each IZ and diffuse IZs in which initiation can occur almost anywhere. However, we can rule out the hypothesis that human replication initiates at isolated, efficient, well-defined replication origins. In particular, we see no evidence for unique initiation sites at the lamin B or Top1 loci (Figure S4A), which have been proposed to be isolated origins (Keller et al., 2002; Abdurashidova et al., 2000). We suspect that the inefficient nature of human IZs and the low signal-to-noise ratio of previous initiation-mapping technology made it difficult to comprehensively map initiation sites across IZs. However, our data suggests that human IZs contain multiple low-efficiency initiation sites, as has been previously suggested (Langley et al., 2016; Petryk et al., 2016; Mesner et al., 2003; Hamlin et al., 2008).

### Human Replication Initiation Events are not Clustered

Our single-molecule data also allows us to directly measure correlations between initiation sites on individual chromosomes. A number of previous reports have claimed that initiations events are clustered, in that initiation events are more likely to occur next to one another on chromosomes than would be expected by chance, and that these clustered events are regularly spaced between 75 and 150 kb apart in human cells (Jackson and Pombo, 1998; Cayrou et al., 2011; Lebofsky et al., 2006; Huberman and Tsai, 1973; Blow et al., 2001; Marheineke and Hyrien, 2004). However, these observations have been largely anecdotal because they have recorded observations only on fibers that have multiple initiation events, and so have not rigorously measured their frequency relative to single initiation events. To test whether initiation events tend to occur in clusters, we examined the frequency and distribution of multiple tracks on single fibers (Figure 3E). If events on a line are clustered with a characteristic inter-event distance, the distribution of inter-event distances will be bell-shaped, with a peak at the most frequent inter-event distance. If, however, the events are distributed randomly, with no inter-event correlation, the distribution of inter-event distances will be exponential (Birnbaum, 1954). We find that, between 60 and 600 kb, the distribution of inter-initiation-event distances is exponential (r = 0.99), inconsistent with any significant clustering of initiation.

As an orthogonal test of initiation clustering, we directly examined co-replication of neighboring IZs. We examined every pair of IZs on each fiber we collected and asked whether the frequency with which they were replicated was correlated (Figure 3F). Using a model that takes into account both passive replication predicted by incorporation track lengths (Figures S2B) and nucleotide-uptake heterogeneity (Figure S1C) but otherwise assumes no correlation among initiation events, we can account for the observed frequency of replication in neighboring IZs (Figure 3F), consistent with the conclusion that human replication initiation sites are not clustered in early S phase. Thus, although initiation events with similar replication timing tend to be clustered because regions of similar replication timing are grouped together (Figure 1F), initiation events are no more likely to occur next to one another, or in collinear clusters, than expected by chance (Figures 3E,F).

### Stochastic Usage of Initiation Zones in Human Cells

The heterogeneous, uncorrelated nature of replication initiation that we observe is reminiscent of stochastic models that have been proposed to explain the regulation of replication timing in budding yeast. Such models propose that the average timing of initiation in a population is regulated by the probability of a site initiating in an individual cell. Sites with a high initiation probability are, on average, more likely to initiate early and thus have an earlier average replication time, although sometimes they initiate late. Sites that have a lower initiation probability are, on average, less likely to initiate early and thus have a later average replication time, although sometimes they initiate early. In particular, neither hierarchical regulation of initiation timing nor correlation between initiation at neighboring sites is necessary in such models. Previous analyses of human replication kinetics have suggested positive correlations, in which one initiation event, or the forks it produces, would increase the probability of other nearby initiation events (Guilbaud et al., 2011; Löb et al., 2016). However, a more recent, higher resolution replication timing study found no evidence of this so-called “domino model” (Zhao et al., 2020). In any case, we find that a stochastic model assuming independent origin initiation can explain our observed data (Figure 3F).

To test whether our results are compatible with a rigorously stochastic model of replication kinetics, we used a computational model of DNA replication that assumes stochastic origin firing (Gindin et al., 2014a). Using our initiation-site mapping data to initialize the model produces predicted replication timing profiles that give an excellent fit (r = 0.85) to experimentally produced profiles (Figure 5B). Since the computational model does not contain any correlation between initiation events, this analysis suggests that any positive or negative correlations between initiation events must be of sufficiently low magnitude or restricted spatial distribution as to be not required to explain observed genome-wide patterns of replication timing. Our work, in combination with work from budding and fission yeast (Kaykov and Nurse, 2015; Yang et al., 2010; Patel et al., 2006; de Moura et al., 2010), supports the hypothesis that stochastic control of initiation timing is a fundamental feature of DNA replication, conserved across eukaryotes.

### General Application of ORM to Map Replication Kinetics

In addition to mapping replication initiation sites, we also used ORM to map ongoing replication forks in unperturbed, asynchronous cells. We inferred fork direction from the asymmetric nature of incorporation tracks caused by the reduction of incorporation as labeled nucleotides are depleted, an approach that has been used before for both radio- and fluoro-labeled nucleotides (Müller et al., 2019; Huberman and Riggs, 1968; Hennion et al., 2020). Averaged over our nearly-500-fold fiber coverage of the genome, our inferred fork directions allow us to infer genome-wide replication kinetics that agree with independently determined replication-timing and replication-fork-directionality profiles (Figures 2C,D). In future work, it should be possible to use sequential labeling with nucleotides of different colors to unambiguously determine fork direction, thereby greatly increasing the resolution and sensitivity of asynchronous ORM. In addition to identifying replication initiation sites throughout the S phase of any cell type, sequential labeling will allow investigation of other important aspects of DNA replication, recombination and repair. We envision ORM being used to measure replication fork speed, replication fork arrests and reversal, and sister-chromatid exchange, under both normal growth and replication stress conditions. Such application will make ORM a central technique for studying DNA replication, DNA repair and genome instability.

### Limitations

The primary limitation of current ORM technology is the *in vivo* pulse-labeling of replication. Other single-fiber DNA-replication-mapping approaches, such as DNA combing (Bensimon et al., 1994) and SMARD (Norio and Schildkraut, 2001), use thymidine analogs to label replicated DNA, allowing kilobase-resolution mapping of replication kinetics in a wide range of cell types and organisms. However, neither the two standard thymidine analogs—BrdU and EdU—are compatible with ORM technology; BrdU because anti-BrdU antibodies only recognize single-stranded DNA, but double-strand DNA is required for fiber stretching in the Bionano optical mapping chips, and EdU because the copper catalyst required to conjugate the EdU to a fluorophore produces hydroxyl radicals, which breaks the DNA fiber into sub-hundred-kb pieces. We have therefore labeled our cells by electroporating them with fluorescently-conjugated dUTP, which limits us to using cell lines. Moreover, the conjugated nucleotides are not efficiently incorporated leading to the sparse labeling and limited resolution that we see. DNA-replication labeling with a number of other nucleoside analogs has been demonstrated recently (Neef and Luedtke, 2014; Rieder and Luedtke, 2014; Mitter et al., 2020). Although none of them are currently compatible with ORM, we predict that improved nucleoside-labeling technology will allow ORM to by applied to most cells, tissues and organisms at kilobase resolution, making ORM a generally applicable technique for replication mapping.

## Methods

### Lead Contact

Further information and requests for resources and reagents should be directed to and will be fulfilled by the Lead Contact, Nick Rhind.

### Materials Availability

No newly generated materials are associated with this paper.

### Data and Code Availability

All raw and processed ORM data, including the identified IZs, is available at figshare <https://doi.org/10.6084/m9.figshare.c.5321048> All ORM analysis code is available at GitHub <https://github.com/cl-chen-lab/ORM>, as are scripts to process ORM data for display on the ORM browser at <https://github.com/ClaireMarchal/ORM2BED> and <https://github.com/nimezhu/ORM>.

### Experimental Model and Subject Details

This study used human HeLa S3 (RRID:CVCL_0058) and H9 cell lines (RRID:CVCL_9773). Both lines are derived from female donors. HeLa S3 cells were grown in DMEM plus 10% Cosmic Calf Serum (GE Life Science SH30087) and Pen/Strep. H9 hESCs were grown in feeder free conditions on Geltrex matrix (Thermo Fisher A14133) coated dishes in StemPro (Thermo Fisher A100701) media according to manufacturer’s specifications.

## Method Details

### Cell Labeling and Sample Preparation

For asynchronous HeLa datasets, cells were grown to ~70% confluency, washed with warm PBS, trypsinized, and electroporated (Lonza, Nucleofection kit SE, HeLa S3 HV program) in the presence of 40 *μ*M Aminoallyl-dUTP-ATTO-647N (Jena Bioscience NU-803-647N). For G1/S synchronized data sets, cells were synchronized at metaphase by incubation with 0.05 *μ*g/ml nocodazole for 4 hours followed by shake off. The percentage metaphase cells was quantified by metaphase spread. Briefly, 3X volumes of ddH20 was added to an aliquot of media from shake off and was incubated at 37°C for 15 minutes. Cells were fixed by adding several drops of ice-cold methanol:glacial acetic acid (3:1). Cells were spun down and pellet were resuspended in ice-cold methanol:glacial acetic acid (3:1) and spotted on to a clean slide. Slides were allowed to dry, spotted with Vectasheild (Vector Laboratories H-1000) containing DAPI and covered with coverslips. Metaphase and interphase cells were counted on a Nikon Eclipse Ti microscope. Metaphase cells were released into fresh media containing 10 *μ*g/ml aphidicolin (Millipore Sigma 178273) for 16 hours. For 0-minute release datasets, cells were washed three times with warm PBS (or PBS plus aphidicolin, in the case of B.0) to remove aphidicolin, trypsinized, and electroporated (Lonza, Nucleofection kit SE, HeLa S3 HV program) in the presence of 40 *μ*M Aminoallyl-dUTP-ATTO-647N (Jena Bioscience NU-803-647N). Cells were recovered in warm media plus aphidicolin for 30 minutes in a 37°C water bath with agitation every 5 minutes to prevent cells from settling in the tube. Cells were then spun down and washed 3 times with warm PBS to remove aphidicolin. For timed release experiments, cells were washed three times with warm PBS to remove aphidicolin, warm media was added, and cells were returned to 37°C/5% CO2 incubator for the indicated times (5, 10, 20, 30, 45, 60, or 90 minutes). Cells were then washed again with PBS, trypsinized, and electroporated in the presence of fluorescent nucleotides as above. In all cases, cells were allowed to recovered overnight in media at 37°C/5% CO2. After recovery, all cells were detached with trypsin and snap frozen in liquid nitrogen.

H9 cells were grown to ~70% confluency, washed with warm PBS, dissociated to single cells with Gentle Cell Dissociation Reagent (StemCell Technologies 07174) and electroporated (Lonza, Nucleofection kit P3, hESC program) in the presence of 40 μM Aminoallyl-dUTP-ATTO-647N (Jena Bioscience NU-803-647N). Cells were recovered overnight in StemPro media plus 10uM Y-27632 dihydrochloride (ROCK inhibitor, Sigma Y0503) at 37°C/5% CO2.

Nucleotide labeling efficiency was assayed by plating a small aliquot of cells on coverslips after electroporation. Following overnight recovery, cells were fixed with 4% formaldehyde for 10 minutes at room temperature. Cells were then washed twice with PBS followed by permeabilization and counterstaining with PBS plus 0.2% Triton X-100 plus DAPI for 5 minutes at room temperature. Coverslips were washed twice with PBS, spotted with Vectasheild, and placed on slides. Images were captured on a DeltaVision (GE Life Sciences) microscope. Replication foci patterns were categorized as previously described (O’Keefe et al., 1992).

### Optical Replication Mapping

Two Bionano mapping approaches were used: Nick, Label, Repair and Stains (NLRS) and Direct Label and Stain (DLS) (Table S1). For NLRS, cell pellets were thawed on ice, resuspended in cold Cell Buffer (Bionano 80004) and embedded into low-melting point agarose plugs using the CHEF Mammalian Genomic DNA Plug Kit (Bio-Rad #1703591) according to the manufacturer’s instructions, with a target concentration of 1,000,000 cells per plug. Cells immobilized in plugs were lysed and treated with Proteinase K and RNase A. The plugs were then melted and solubilized, and the resulting DNA was further cleaned by drop dialysis. The DNA was fluorescently labeled with green fluorophores at Nt.BspQI sites and counterstained with YOYO-1 using Bionano Genomics’ NLRS labeling kit (Bionano 80001). Throughout the entire process, the samples were shielded from light whenever possible, in order to maintain integrity of the fluorophores. For DLS, DNA was labeled with green fluorophores using the direct labeling enzyme DLE-1 (Bionano Genomics 80005) according to manufacture’s protocol. The rest of processing steps were the same as NLRS labeling.

Labeled DNA samples were loaded onto a Saphyr chip (Bionano Genomics 20319) and run on the corresponding Saphyr instrument (Bionano Genomics IN-011-01), following the manufacturer’s instructions. Briefly, the DNA was coaxed into an array of parallel nanochannels via electrophoresis, thereby elongating the DNA to a uniform contour length, allowing for accurate measurement of distances along the molecules. Molecules were imaged to collect the YOYO-1 DNA signal in a false-color blue channel, the Nt.BspQI or DLE-1 labels in the green channel, and the *in vivo* replication incorporation tracks in the red channel.

Images were converted to digitized molecules and the positions of the green and red labels on each fiber were determined using Bionano Access. The Nt.BspQI and DLE-1 labels were aligned to human reference genome hg19 by creating an *in silico* digestion of the reference FASTA file. Alignment of the fibers’ Nt.BspQI and DLE-1 sites allowed mapping of the red replications signal to the genome. The pipeline produces one .bnx file containing the intensity and location of each green and red signal on each fiber and one .xmap file containing the genomic location of each fiber. We observed abnormal ORM signal enrichment in some specific genomic positions (hereafter called hotspots), which only occurred in the samples mapped by NLRS technique but not those mapped by DLS technique. Further investigation confirmed that most (99.36%) these hotspots are located within < 650 bp distance to a BspQI site used for the mapping in the NLRS technique. Detailed analysis showed that these abnormal ORM signals are likely signals of false positive detection due to the strong BspQI signals, since the green signals (for mapping) associated with the hotspot regions have stronger intensity distribution than the green signal outside the hotspot region while the red signals inside the hotspot regions have intensity weaker than the red signals outside of the hotspots. In order to filter these false positive ORM signals, we calculated the ORM signal count (mean = 1.71 and s.d. = 6.35 for all windows containing at least one red signal) within all 300 bp windows by combing all 0’ data, and removed those with >=20 ORM signals (the cutoff was estimated by comparing with the distribution of DLS samples without such hotspots). The removed hotspot signals correspond to 4.7% of total signals in C.0 sample and 3.6% in D.0 sample, and to similar degree (~3-4%) for the other samples from these datasets (i.e. C.5, C.10, D.20, D.30, D.45, D.60 and D.90).

### Modeling Signal-Intensity and Inter-Signal-Distance Distributions

To model the signal-intensity distributions (Figure S2A), we assumed that signals represent individual fluorophores that can be resolved with some finite resolution. We modeled both the photon output and the inter-signal distance (which controls the probability of having two fluors closer together than the resolution of the Saphyr) of each fluor as Poisson processes, resulting in the depicted fit (obtained from the first four terms of Eq. 5, Supplemental Mathematical Methods), which predicts that 20% of the fluors are closer than the resolution of the Saphyr. See Supplemental Mathematical Methods for details.

To model the distribution of inter-signal distances (Figure S2B), we assume that the incorporation rate is a Poisson process for which the Poisson statistic is determined by the average amount of label taken up by cells. We further assume that there is an exponentially decreasing amount of label over time as it is consumed by replication. Combining these two assumptions results in the depicted fit (Eq. 14, Supplemental Mathematical Methods), with an inferred initial incorporation rate of about 1/900 thymidines and a depletion half length of about 75 kb for the merged 0-minute dataset. See Supplemental Mathematical Methods for details.

Combining the models in Figures S2A and B, we calculate that 20% of the signals in Figure S2B (corresponding to the 20% of the fluors that we infer from Figure S2A are not individually resolved by the Saphyr) are located within 1.3±0.3 kb of a neighboring fluor, consistent with the reported resolution of the Saphyr and the observation that there are almost no inter-signal distances below 1.3 kb in our datasets.

### Identification of Replication Tracks

After mapping the fibers on the human genome, the distances between the genomic mapping signals were recalibrated to match their reference genome positions, compensating for heterogeneity in fiber stretching. We then mapped signals in the red channel between two adjacent genomic mapping signals as nucleotide-incorporation signals. The nucleotide-incorporation signal is discontinuous and appears as clusters of neighboring signals (Figure 1A). To segment the ORM signals into replication-incorporation tracks, we used a two-step process based on Gaussian mixture models. We first fit a Gaussian mixture model using the R package *mixtools* (https://www.rdocumentation.org/packages/mixtools) with 3-gaussian distributions to the inter-signal distance distribution, with the first peak corresponding to the distances between signals within the same segment, the second peak corresponding to the distances between signals in adjacent segments and the third peak corresponding to signals in non-adjacent segments. The tail of the first Gaussian distribution (16,384 kb) was comparable amongst various data sets and was therefore selected as the cutoff value (cutoff1) to merge adjacent signals into primary segments. We then measured the distance between adjacent primary segments and fit a 2-Gaussian distribution to find the second cutoff value (cutoff2, 32,768 kb), which we used to merge all the adjacent primary segments into final replication tracks.

To estimate the resolution with which we can identify the initiation site of any given replication track, we defined the maximum likelihood distribution of initiation sites for a given replication track and, using numeric simulations, estimated the standard deviation of that distribution as 14.2±0.1 kb. See Supplemental Mathematical Methods for details.

To determine the direction of replication tracks in our asynchronous data, we defined a fork direction index (FDI): FDI = log2(IntensitySUM_Left / IntensitySUM_Right), where IntensitySUM_Left and IntensitySUM_Right is the total fluorescent signal in the left and right half, respectively, of each ORM track with at least 3 signals. A positive (negative) FDI corresponds to a rightward (leftward) moving replication fork. The mean relative fork direction (RFD) within a given window is defined as (R-L)/(R+L), where R and L is the number of rightward (FDI>0) and leftward (FDI<0) ORM tracks, respectively.

### Identification of Initiation Zones

Initiation zones were identified based on the ORM signal density along the genome. We first obtained the ORM signal density by calculating the percentage of fibers containing ORM signal in 10 kb sliding windows with 1 kb steps. LOESS smoothing (with α = 0.75 and the polynomial degree = 2) was then performed in successive 160 kb windows, with 8 kb overlaps to avoid discontinuity of fitting. Within the 8 kb overlaps, the final smoothed values were the averages, weighted by their distance to the corresponding window, of the smoothed values from two adjacent windows. IZs were then defined as the peaks on the smoothed ORM signal density profile. The size of each IZs was defined by the smallest window that containing at least 40% of ORM signals within each peak. The value of 40% was chosen because, when aggregated around IZ peaks, the ORM signal density decreases with distance from the peak at a similar rate until about 40% then decreases much more slowly. This process was performed on all 4 biological replicates of HeLa 0’ data as well as the combined dataset. Only IZs identified in at least 3 replicates and with a relative peak height in the combined dataset (as measured by the difference in signal density between the peak and the edges of the zone) of greater than 0.3% were retained. The precise location of each retained IZ was then determined by analyzing the transition of ORM signals with a K-mean clustering (n=2 if the center cluster < 30 kb, otherwise n=3) on the ΔORM signal between adjacent 1kb windows to identify the region with sharpest ORM density around each IZ center.

To estimate the minimum number of initiation sites in each IZ, we calculated number of replication track centers separated by more than the resolution of our ability to localize initiation sites, which is about 15 kb. We calculated the position of the center of each replication track that overlaps a given IZ, discarding any that are more than 7.5 kb outside of it. We then ranked the centers by position and, starting at one end, counted the number of centers that are more than 15 kb apart.

### Comparative Analysis of ORM Data with Other Datasets

The raw Repli-Seq data of HeLa S3 cells were downloaded from the Encode project <http://genome.ucsc.edu/cgi-bin/hgFileUi?db=hg19&g=wgEncodeUwRepliSeq> (Hansen et al., 2010) and S50 (the fraction of S phase at which 50% of the DNA is replicated in a defined genome region) was computed (Chen et al., 2010) and used to identify replication timing peaks (Chen et al., 2011) as previous described. Published H9 ESCs replication timing and peak calling data were used (Zhao et al., 2020). ORM signal enrichment around timing peaks was calculated in R (version 3.5.1 <https://www.r-project.org>). To compare the ORM IZs and other origin mapping data, we used pROC package to calculate the ROC (Receiver Operating Characteristic) curve and AUC (Area Under the Curve) values by using all 100 kb bins along the genome as example space and the bins overlapped with the ORM IZs as true positive space (Robin et al., 2011).

Correlations between ORM initiation efficiency and published genome-wide HeLa cell replication-initiation-mapping datasets—OK-seq (Petryk et al., 2016), SNS-seq (Picard et al., 2014), Ini-seq (Langley et al., 2016) and Orc1 ChIP-seq (Dellino et al., 2013)—were calculated across the genome in 50 kb adjacent windows. When necessary, genomic coordinates were remapped to hg19 using LiftOver (https://genome.ucsc.edu/cgi-bin/hgLiftOver). ORM initiation efficiency was calculated as the number of fibers with ORM signal in a 50 kb genomic window divided by the number of fibers covering that window. The OK-seq initiation efficiency was calculated either genome-wide using the previously described origin efficiency metric (OEM) approach (McGuffee et al., 2013) or for each OK-Seq IZ with the ΔRFD between its right and left extremities (Petryk et al., 2016). Correlations were calculated using Spearman’s correlation coefficient in R.

The genome-segmentation results (ChromHMM) based on ENCODE data were retrieved from UCSC genome browser <https://genome.ucsc.edu/cgi-bin/hgFileUi?db=hg19&g=wgEncodeAwgSegmentation>. To compensate for the large size of IZ, relative to many of the ChromHMM segments, we used the IZ’s core regions, defined as the region from center of the IZ to the point of highest ORM signal within that IZ.

Histone modification data for HeLa S3—H2A.Z, H3K4me1, H3K4me2, H3K4me3, H3K9ac, H3K9me3, H3K27ac, H3K27me3, H3K36me3, H3K79me2 and H4K20me1—was downloaded from ENCODE <https://www.encodeproject.org>. If available, the data was obtained from G1 phase cells. The average of all replicates was use in our analysis. The RNA Pol II-pS2 <https://www.encodeproject.org/experiments/ENCSR000ECT> and DNase I hypersensitivity data (UW DNase I HS data track) for HeLa S3 were also downloaded from ENCODE. The signal enrichment of each feature was re-normalized such that the background regions show zero log2 enrichment ratio. Published G4 sequence locations were retrieved (Puig Lombardi et al., 2019). We analyzed 3 G4 datasets (G4L3, G4L7 and G4L12) with similar results; we therefore used the G4L12 (G_3_N_1-12_G3N_1-12_G3N_1-12_G_3_), which contains the largest dataset and give the best correlation with ORM data, in our final analysis. The genome.wide pairwise correlation analysis and enrichment around ORM IZs were performed by deepTools (Ramírez et al., 2016) or with R (version 3.5.1 <https://www.r-project.org>). IZs were classified by timing—early (S50<0.25), mid-early (0.25 ~ 0.5), mid-late (0.5~0.75) and late (S50>0.75)—using the S50 of the IZ center. The partial correlation analysis between ORM initiation efficiency and G4 sequences with controlling the effect of GC percentage (obtained from UCSC <http://hgdownload.soe.ucsc.edu/goldenPath/hg19/gc5Base>) was performed by using the R package ppcor <https://cran.r-project.org/web/packages/ppcor/index.html>. Features were visualized in IGV (Robinson et al., 2011) and the ORM Browser <http://orm.nucleome.org> (Figure S3, Zhu et al. in prep, personal communication, J. Ma <https://github.com/nimezhu/ORM>). All published data sources are complied in Table S2.

### Correlation of Initiation in Neighboring Initiation Zones

To calculate the degree of correlation between initiation events in neighboring IZs, we defined an initiation event as any ORM signal in any IZ on a fiber. We then calculated the correlation as the frequency with which we observed initiation in neighboring IZs on the same fiber divided by the product of the frequency of overall initiation in the two IZs. The non-neighbor correlation was calculated in the same manner, but using all pairwise combinations of IZs on each fiber farther that 200 kb apart. To calculate the expected correlation due to passive replication of one IZ due to initiation in a neighboring IZ, we used the observed label-depletion rate and inferred the frequency with which initiation at one IZ would lead to incorporation of label at a neighboring IZ at a given distance away (Figure S2B). To calculate the correlation expected from heterogeneity of label uptake by cells, we used the observed distribution of cellular labeling (Figure S1C). The two calculations were combined to produce the model (Eq. 34, Supplemental Mathematical Methods) shown in Figure 3F. See Supplemental Mathematical Methods for details.

### Replication Timing Simulation

To simulate the replication timing profiles, we used the Replicon simulation code (Gindin et al., 2014a). Replicon uses three sets of user-defined parameters. The first set of parameters is the initiation probability landscape (IPLS), the relative probability of initiating at any point in the genome. In our simulations, we used the ORM the signal distribution or other genomic features with a bin size of 0.5 kb. The second set of parameters control the virtual flow sorter. In agreement with the results from the original paper (Gindin et al., 2014b), we set this to (0.0, 0.17, 0.35, 0.58, 0.92, 0.99, 1.0), although other choices give similar results. The final parameter is the number of forks. Here, we again followed the original paper and set the number of forks equal to 10.24 + x(7.9E-7), where x is the length of the chromosome in base pairs.

### Quantification and Statistical Analysis

Statistical analyses were performed in R. Statistical tests and parameters are reported in the text and figure legends.

### Additional Resources

Interactive visualization of the ORM data is available on the ORM Browser <http://orm.nucleome.org>.

## Acknowledgements

We are grateful to Feng Yue for his contribution to the collection of the Bionano data, Jian Ma for help developing ORM browser and Peiyao Zhao maintaining the ORM browser server. The work was funded by NIH grants HG010658 to DMG and GM125872 to NR. CLC was supported by the I. Curie YPI program, the ATIP-Avenir program from CNRS, Plan Cancer from INSERM, the CNRS 80|Prime interdisciplinary program, ANR and INCa. WW was supported by a COFUND IC-3i International PhD fellowship.

## Author Contributions

**Conceptualization:** CLC, DG, NR, **Methodology:** WW, KK, KP, AH, JB, CLC, DG, NR, **Investigation:** WW, KK, KP, HY, **Software:** WW, KP, MC, XZ, CLC **Validation:** WW, KK, KP, AH, HY, JB, CLC, DG, NR, **Formal Analysis:** WW, KP, JB, CLC, NR, **Resources:** AH, CLC, DG, NR, **Data Curation:** WW, CLC, **Writing – Original Draft:** NR, **Writing – Review & Editing:** WW, KK, KP, AH, HY, JB, CLC, DG, NR, **Visualization:** WW, KP, CLC, NR, **Supervision:** JB, CLC, DG, NR, **Project Administration:** NR, **Funding Acquisition:** AH, CLC, DG, NR

---

**Figure S1.**
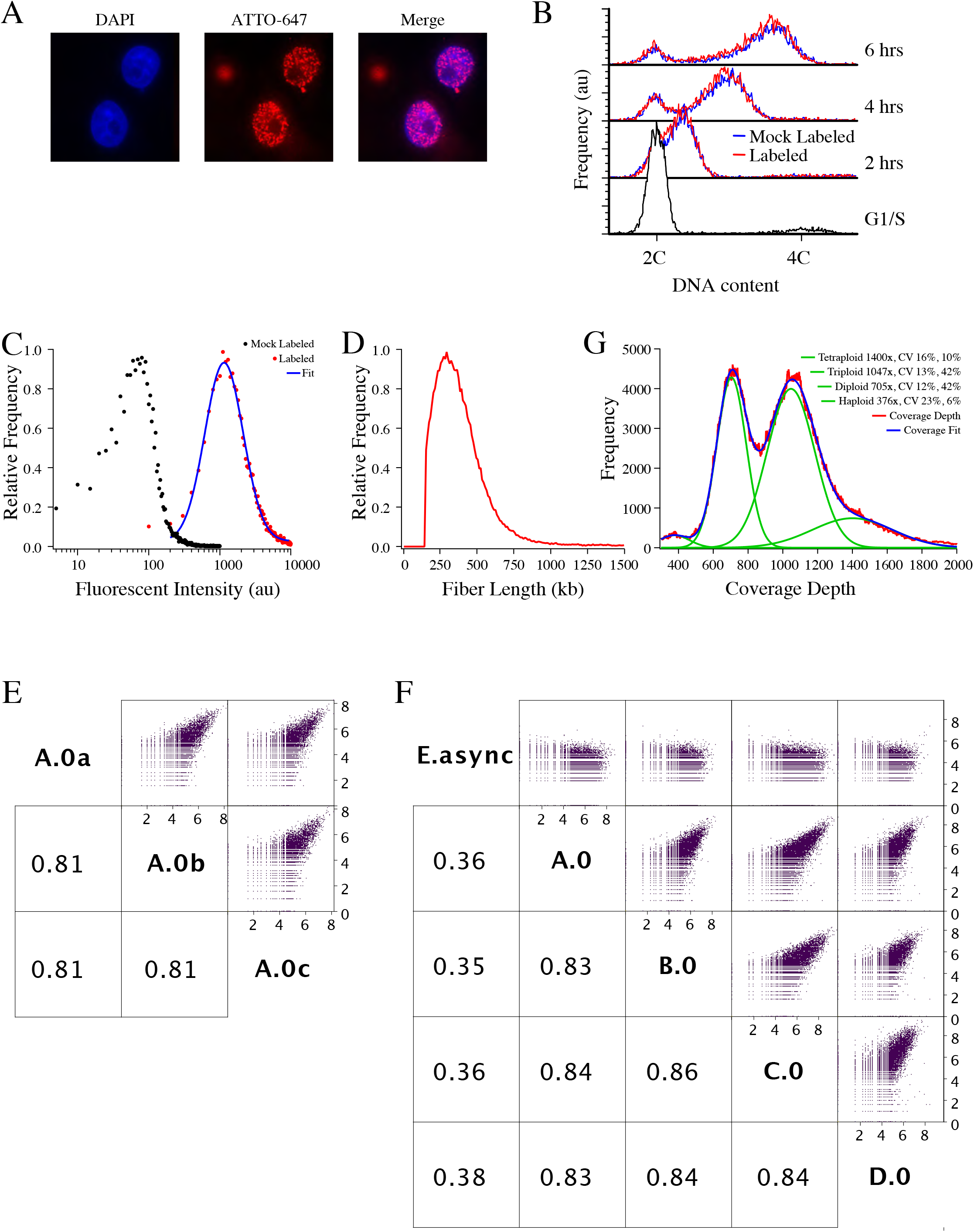
Characterization of ORM Labeling. **A**) Micrographs of representative HeLa cells electroporated with ATTO-647-dUTP during an aphidicolin arrest, released, allowed to recover overnight, and fixed. The left panel is stained with DAPI, the middle panel visualizes the incorporated fluorescent nucleotide and the right panel is a merger of the two channels. **B**) Flow cytometry analysis of S-phase progression after ATTO-647-dUTP electroporation. **C**) Flow cytometry analysis of ATTO-647-dUTP uptake. Cells arrested in aphidicolin at the beginning of S phase were electroporated with ATTO-647-dUTP, or mock electroporated, incubated on ice and analyzed by flow cytometry. The distribution of labeled cells was fit with a log-normal distribution with a mean of 1141 ± 7.7and a coefficient of variation of 0.88 ± 0.01. **D**) The distribution of fiber lengths of 0-minute dataset B.0, which was collected with the DLS mapping approach and has slightly longer fibers that datasets collected with the NLRS approach (Table S1). The average fiber length is 338 kb and the fiber N50 is 369 kb. **E**) The depth of genome coverage in the combined 0-minute dataset in 1 kb bins. The aneuploid character of the HeLa genome is evident in the distribution of coverage into four peaks corresponding to the haploid, diploid, triploid and tetraploid regions of the genome. The coverage data was fit with four Gaussian curves with coverage maxima at about 376x (haploid), 705x (diploid), 1047x (triploid) and 1400x (tetraploid, which was fixed at 1400x, because the unconstrained fit had a very large variation). The individual Gaussians and the complete fit are shown. The coefficients of variation of the individual Gaussians and the percent of the genome inferred to have that ploidy are shown in the legend. F) The correlation of labeling between the three biological replicates that make up the C.0 0-minute dataset. The number of signals in each 10 kb bin across the genome is plotted and the correlation coefficient is reported. **G**) The correlation of labeling between the four biological replicates (A.0, B.0, C.0, D.0) of the 0-minute dataset and one asynchronous dataset, as in panel D. The correlations between the biological replicates are higher than those between the technical replicates because the biological replicates are larger, reducing the counting noise in infrequently labeled regions of the genome.

**Figure S2.**
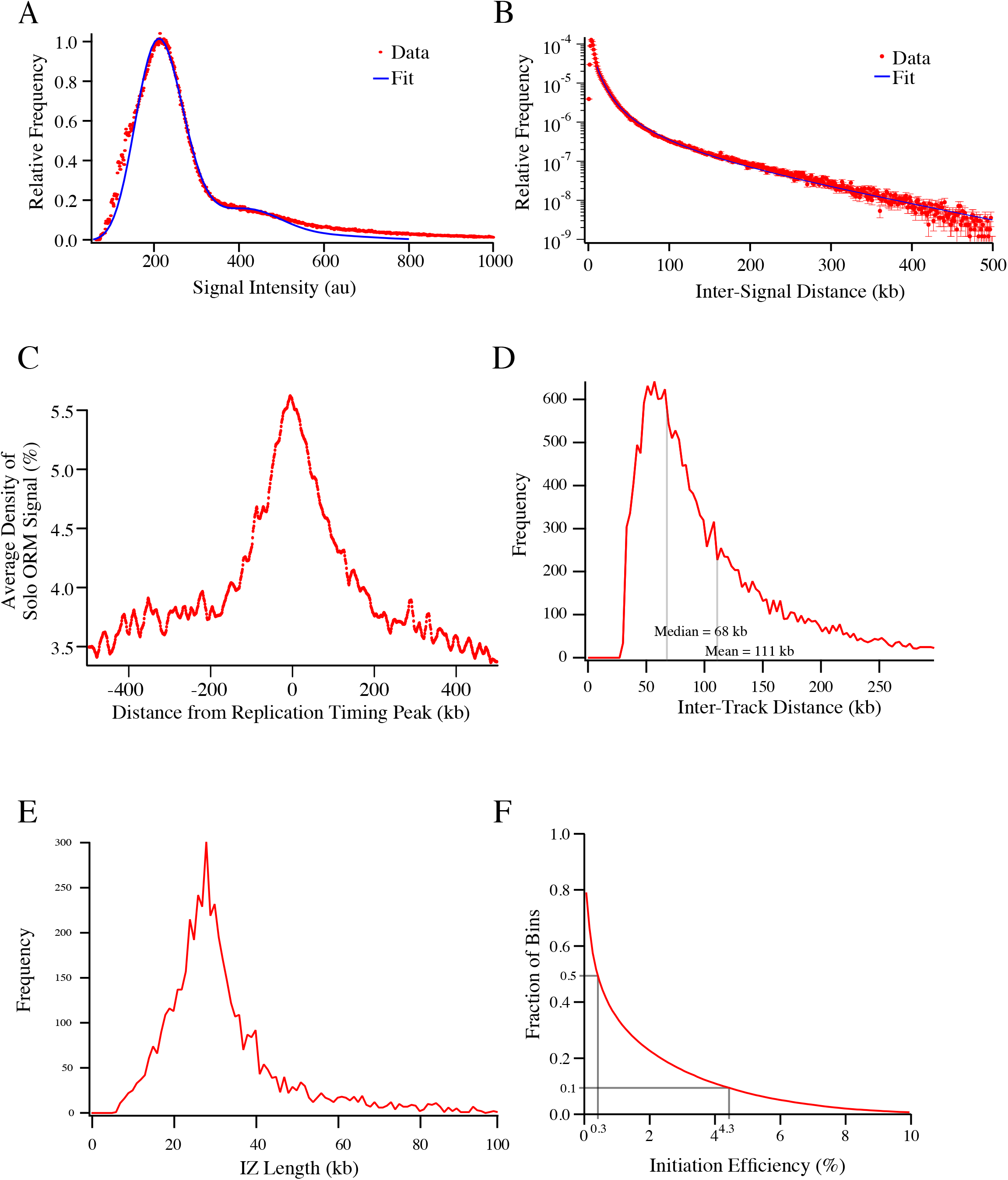
Characterization of ORM Label Distribution. **A**) The distribution of the intensities of the incorporated fluorescence signals in the D.0 0-minute dataset. The fit (obtained from the first four terms of Eq. 5, Supplemental Mathematical Methods) predicts that about 80% of observed signals are single fluorophores and that the other 20% are multiple fluorophores sufficiently close together that they are not resolved by the Saphyr optics. This estimate of 80% single fluorophores is consistent with an average inter-signal distance of 4 kb and 1.3 kb resolution of the Saphyr, both of which parameters can be inferred from the distribution of inter-signal distances (see panel B and Methods). The distribution of intensities in the other datasets are similar, although differences in the Saphyr optical calibration on different runs introduces variation into the absolute value of the measured fluorescent intensities. **B**) The distribution of inter-signal distances in the combined 0-minute dataset. The fit to the data (Eq. 14, Supplemental Mathematical Methods) between 10 and 500 kb predicts an initial labeling frequency of 1 in every 877±17 thymidines and a depletion half length of 74.5±0.7 kb. Similar fits for the asynchronous HeLa and H9 datasets predict labeling frequencies of 1/1025 and 1/850 and depletion half lengths of 57 and 48 kb, respectively. The similar labeling densities suggest the nucleotide uptake is similar in all three experiments, whereas the shorter depletion half length is consistent with previous reports that the number of forks increases during S phase, which would consume nucleotides more quickly (Yang and Bechhoefer, 2008; Goldar et al., 2009). **C**) The enrichment of ORM signals in solo-signal replication tracks from the combined 0-minute dataset around early replication-timing peaks, those that replicate in the first quarter of S phase. **D**) The distribution of inter-replication-track lengths. **E**) The distribution of IZ lengths. **F**) The fraction of 50 kb genomic bin with initiation efficiency greater than indicated on the x axis. 50% of bins have an initiation efficiency greater that 0.3% and 10% of bins have an initiation efficiency of greater that 4.3%.

**Figure S3.**
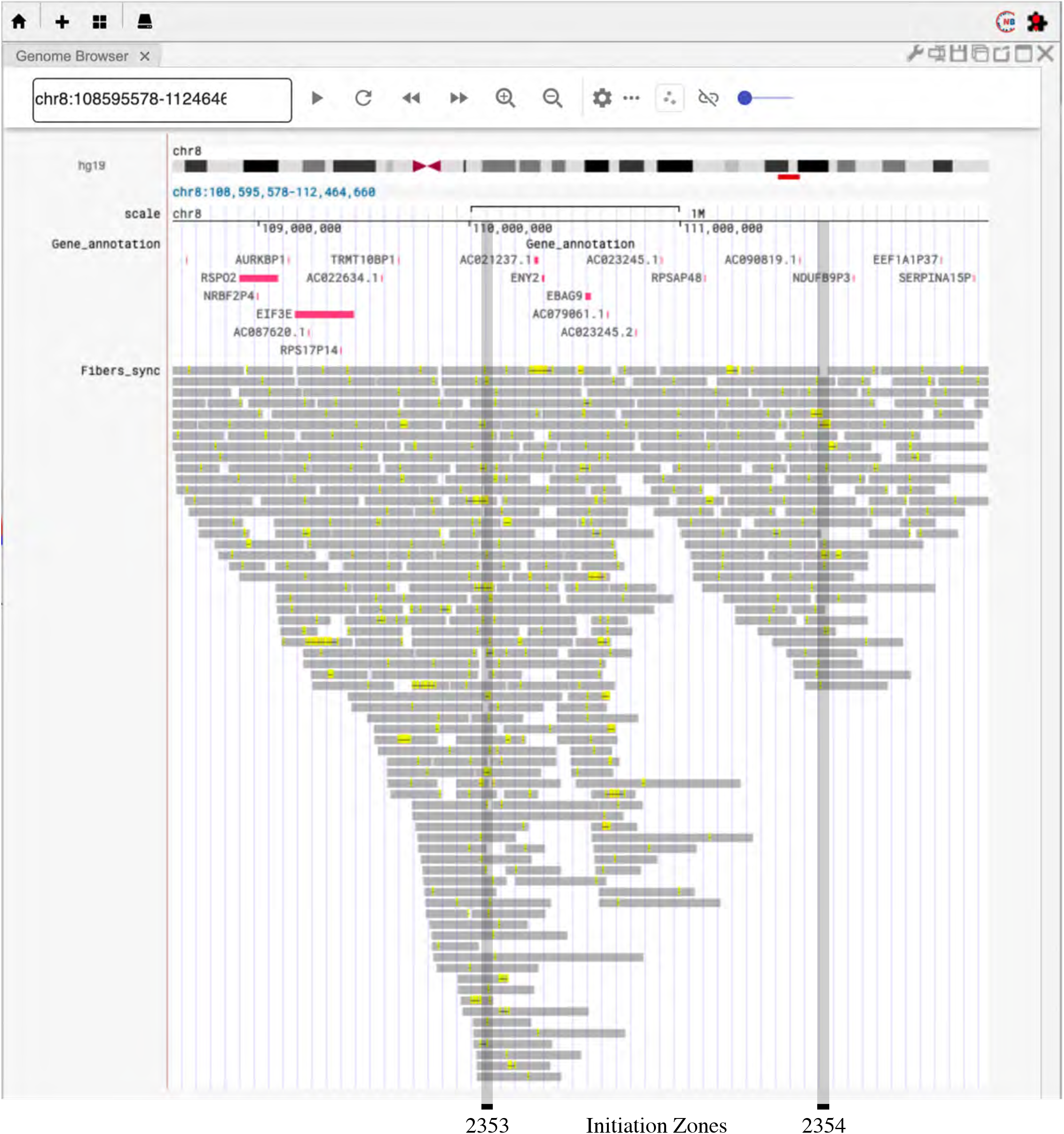
The ORM Genome Browser. The ORM Genome Browser allows interactive visualization the HeLa ORM data. Shown is a screen shot of the Fibers track of synchronous data. It shows fibers as gray bars, ORM signals as yellow hash marks, and inferred replication tracks as black lines. Only fibers labeled with ORM signals are displayed because only~5% of fibers are labeled; displaying all fibers is impractical. The browser can also display only the replication tracks, to provide a more easily-visualized view of replication initiation. It can also show the fibers and the replication tracks from the asynchronous data. Two low-efficiency initiation zones are shown, 2353 (3%) and 2354 (2%), because higher-efficiency initiation zones contain too many labeled fibers. Note that, although the signals and replication tracks are concentrated around the initiation zones, many lay outside of them and none are concentrated in discrete areas.

**Figure S4.**
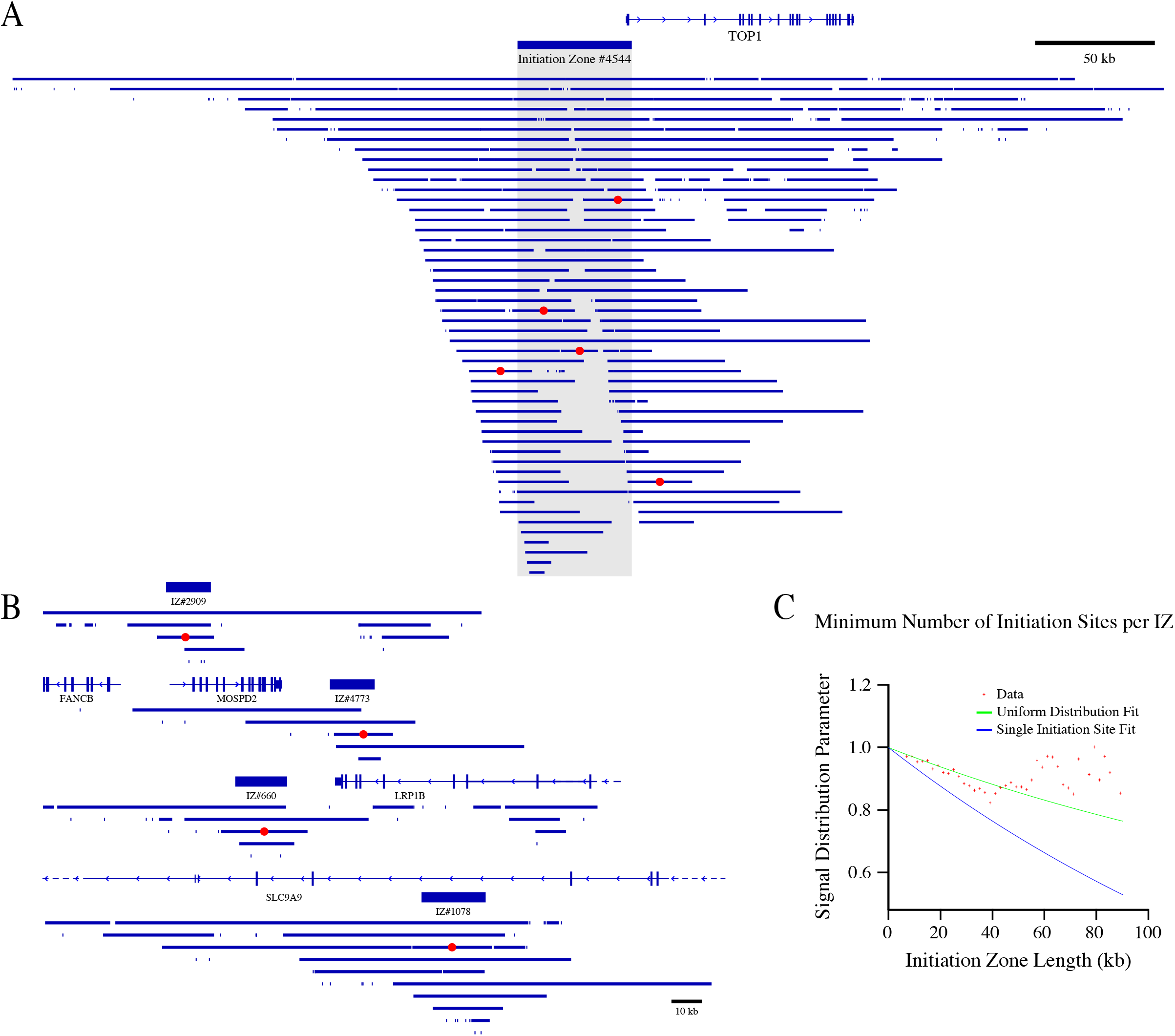
Distribution of Replication Tracks within Initiation Zones. **A**) The distribution of replication tracks in the merged 0 minute dataset at the Top1 locus. The Top1 IZ has an estimated minimum of five initiation sites because the five replication track centers indicated in red are all 15 kb away from each other. **B**) The distribution of replication tracks at four examples of IZs for which our estimate of the minimum number of initiation sites is 1. **C**) The distribution of signal across IZs. The ratio of signal frequency at the IZ center to the IZ boundary is plotted versus IZ length. This value is expected to decrease more quickly in IZs that predominantly have a single initiation site (Eq. 25) than if initiation is distributed across the IZ (Eq. 28, Supplemental Mathematical Methods). The distribution across IZs shorter than 55 kb is consistent with a uniform distribution of initiation sites. At longer lengths, we actually see more signal at the edges of the IZs than it the center. One possible explanation for this phenomenon is that larger IZs may actually be two smaller IZs fused together such that there is more initiation sites towards the edges of the fused IZ and less initiation in the center, which is actually between the two constituent IZs. See Supplemental Mathematical Methods for details.

**Figure S5.**
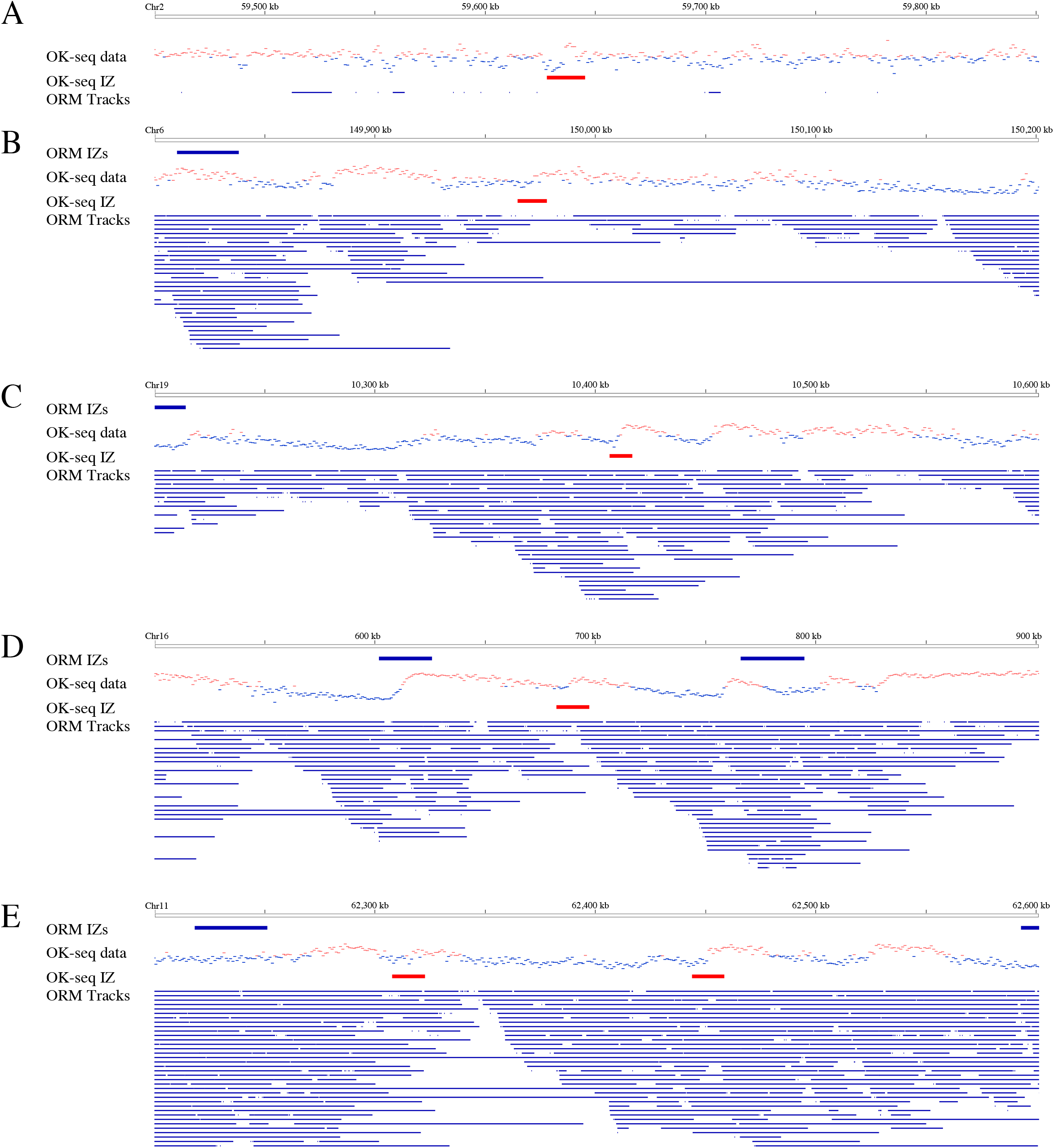
Reanalysis of Potentially Discrete OK-Seq Initiation Zones. We reexamined the 66 OK-seq IZs that were reported to be less that 5 kb wide (Petryk et al., 2016). **A,B**) 53 are in regions of noisy OK-seq data. Of those, 24 are in late-replicating regions and appear to be in regions with extensive bi-direction replication. Panel A is an example of one such zone. However, since there is little ORM data in these regions, we can say little more about them. 29 are in early replicating regions, but none of them correlate with numerous ORM segments. Panel B is an example of one such zone. We conclude that they are not active IZs in our ORM data and probably not active IZs in the OK-seq data, either. **C**) 6 are robust transitions that correlate with numerous ORM segments. We conclude that they are IZs in both the OK-seq and ORM data. However, they show broadly dispersed ORM segments, therefore we do not believe they are unusually constrained IZs. Instead, we conclude that they are outliers in the OK-seq data that were identified as unusually narrow due to experimental variation. Panel C is an example of one such zone. **D,E**) 7 are robust transitions that do not correlate with numerous ORM segments. They could be IZs present in the OK-seq HeLa cell line, but absent in the ORM HeLa cell line. Alternatively, they could be translocation break points in the OK-seq HeLa cell line relative to the hg19 reference sequence. Such breakpoints would explain both the sharpness of the transition and the absence of these putative IZs from the ORM data. Panel D is an example of one such zone. Panel E show two such zones that can be explained by an inversion between them.

**Figure S6.**
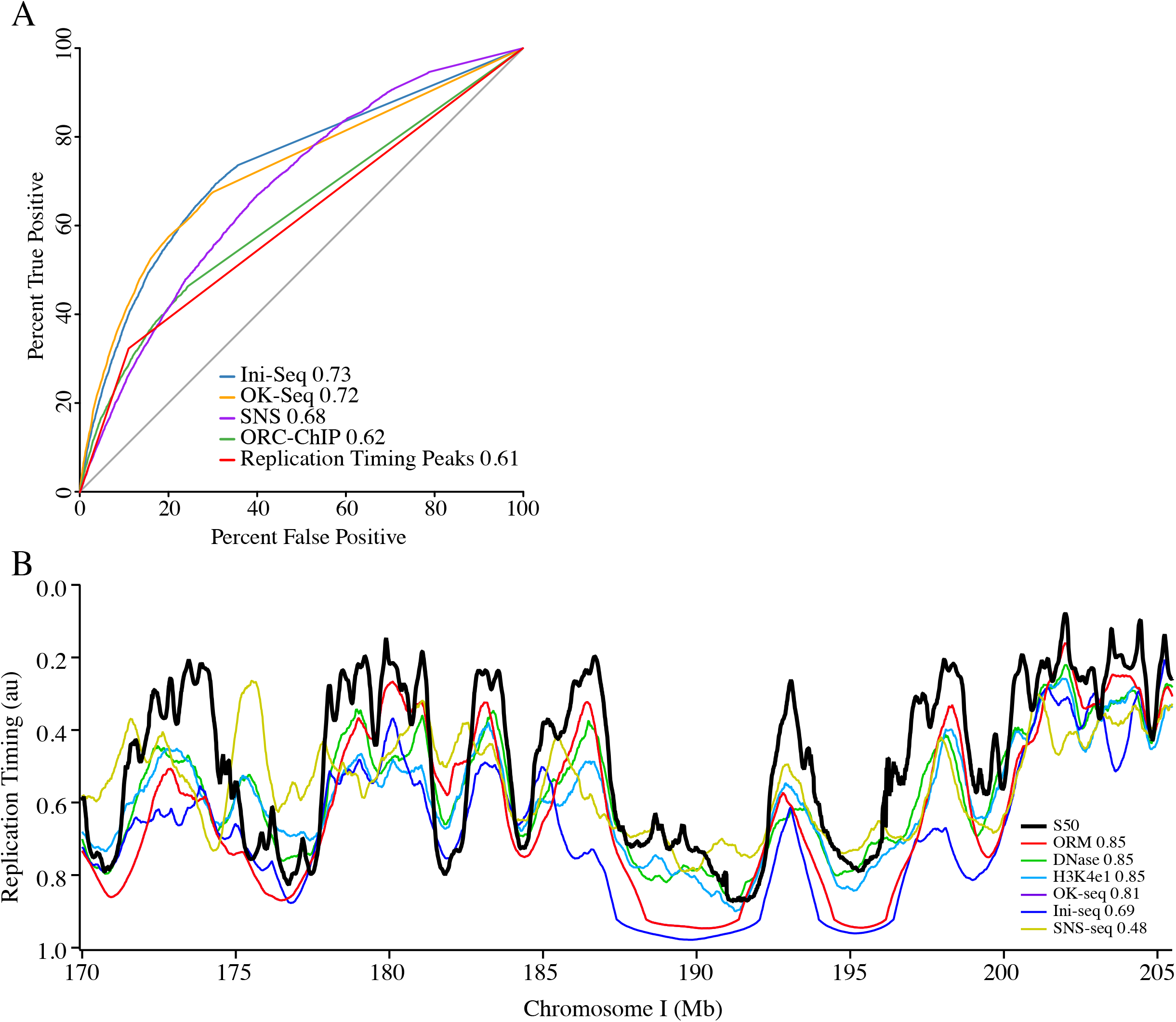
Simulated Replication Timing Profiles. A) ROC analysis of the association between the ORM IZs and other origin mapping data shown in Figure 3G. As shown in the ROC curves, the ORM IZs are better correlated with OK-seq IZs (AUC=0.72) and Ini-seq IZs (AUC=0.73). The bias of the SNS ROC curve towards high true positive rates only at high false negative values is consistent with that dataset having more false positive signal, whereas the bias of the replication timing and ORC datasets relatively high true positive rates only at low false negative values is consistent with those datasets having fewer, but more accurate true positives. Areas under the ROC curves (AUC) are shown in the legend. B) Comparison between experimentally determined HeLa replication timing (S50) and replication timing predicted from ORM, DNase I hypersensitivity (Bernstein et al., 2012), OK-Seq (Petryk et al., 2016), Ini-seq (Langley et al., 2016) and SNS-seq (Picard et al., 2014) data using a stochastic model (Gindin et al., 2014a). The Spearman correlation coefficients with replication timing are shown in the legend.

**Figure S7.**
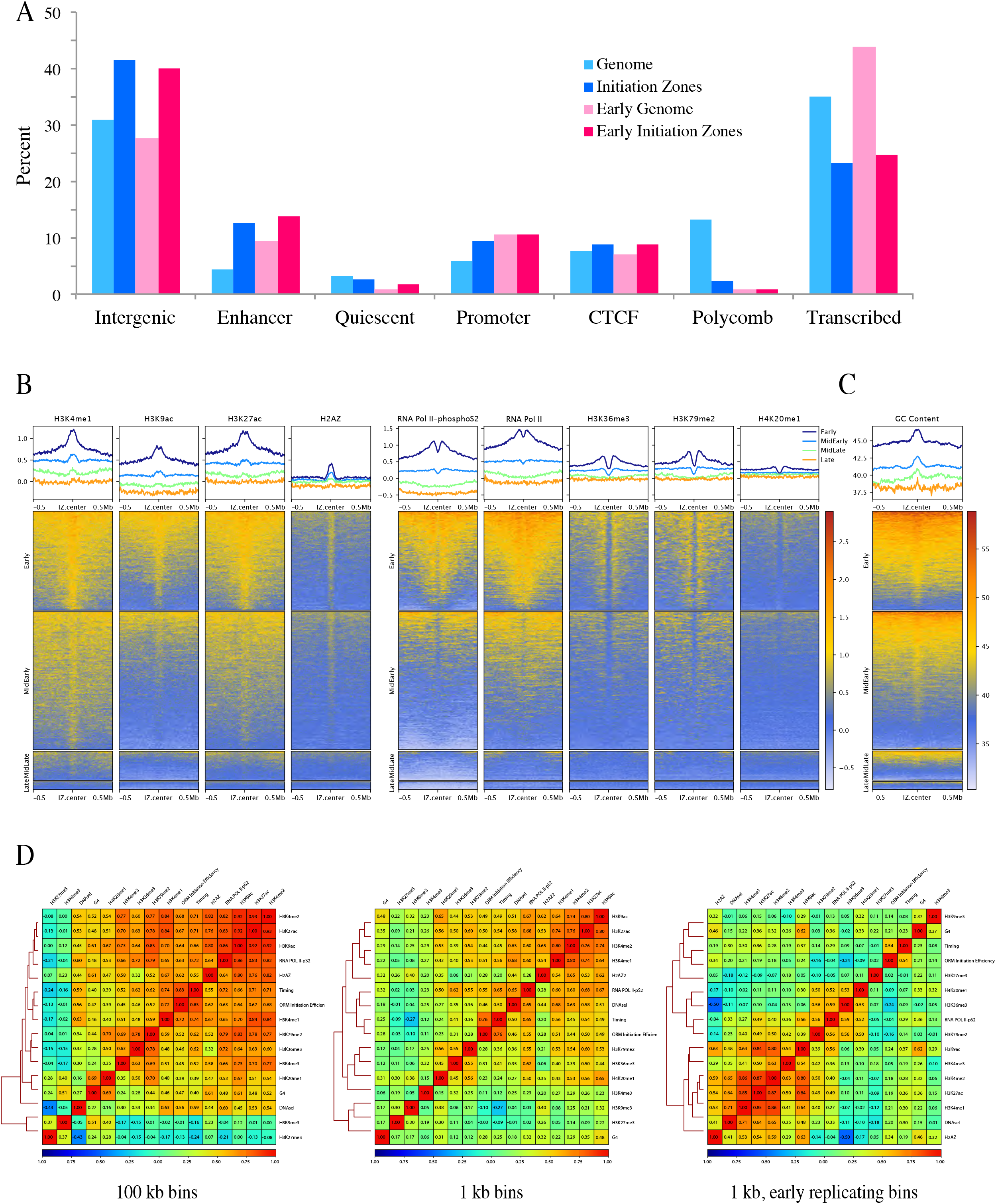
Enrichment of Histone Modifications and Other Genomic Features in Initiation Zones. **A**) The enrichment of chromatin regions defined by ChromHMM in IZs relative to the genome in general (Ernst and Kellis, 2012). The faction of IZs or genomic sequence with each of the indicated annotation is shown for all IZs and the IZs that replicate in the first quarter of S phase (S50 < 0.25). **B**) The enrichment of histone modifications and other genomic features relative to ORM IZs. The upper panels show the genome-normalized relative ChIP-seq signal around all IZs. The lower panels show heat maps of the same signal at each IZ. The left panel includes enhancer-enrich features; the right panel shows transcription-enriched features. **C**) The enrichment of GC content relative to ORM IZs. The upper panels show the average % GC content signal around all IZs. The lower panels show heat maps of the % GC content at each IZ. **D**) Correlation heat maps at various resolution. The left panel shows 100 kb resolution, which does not resolve enhancers, promoters and transcription units. Therefore, features associated with all three correlate with ORM signal. The center panel shows 1 kb resolution, which resolves enhancers, promoters and transcription units. However, the correlation is dominated by replication timing, creating a correlation between ORM IZs and transcription units, which both tend to replicate early. The right panel shows 1 kb resolution for the earliest-replicating quarter of the genome. Here, enhancer-enriched features, such as H3K4me1, H3K9ac and H3K27ac hypersensitivity, are most strongly correlated, while promoter-enriched features, such as RNA Pol II, H3K4me3, and are more weakly correlated and elongation-enriched features, such as H4K20me1, H3K79me2 and H3K36me3, are anti-correlated.

## Supplemental Mathematical Methods

### 1 Modeling the Signal-Intensity Distribution

The intensity of a signal is directly proportional to the number, *n*, of detected photons. Its probability distribution *p*(*n*) results from a combination of two processes: the number of photons coming from each fluorophore and the number of fluorophores inside each resolution-limited region measured. If we assume that the incorporation of fluorophores happens independently, both of these processes are Poisson distributed. The number of photons coming from each fluorophore is Poisson distributed with (unknown) parameter *λ_p_*. Therefore, if there are *N* fluorophores in the measured region, the number of photons is Poisson distributed with parameter *Nλ_p_*. On the other hand, the number of fluorophores N is Poisson distributed with parameter Λ_*f*_. Therefore, the distribution of the number of photons is given by

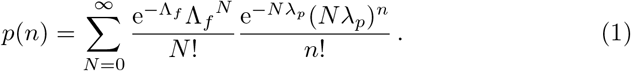

One can simplify this expression as one expects the number of photons (and therefore *λ_p_*) to be large. Therefore, we can use the Stirling approximation [1],

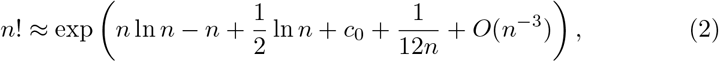

where co is a constant, to rewrite this to

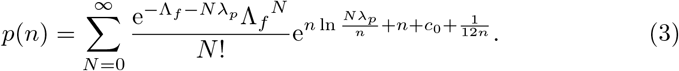

The signal intensity *x* is proportional to the number of photons, *x* = *cn*, with an unknown proportionality coefficient,

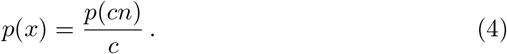

In experimental data, one cannot determine *p*(*x*) for small *x* due to background signals. Therefore, we need to add a renormalisation constant, *a*, in which we can absorb the prefactor exp(−Λ_*f*_ + *c*_0_), to get

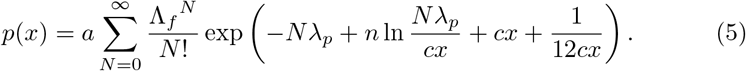

We now have four unknown parameters: *a*, Λ_*f*_, *λ_p_* and *c*. These were found via a fit using gnuplot’s standard fitting procedure (http://www.gnuplot.info), which gives

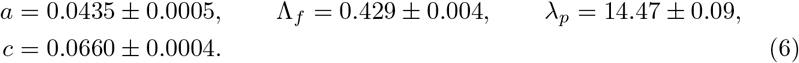

### 2 Probability Distribution of Intersignal Distances

We seek the intersignal distance distribution, *p_ℓ_*(*f*). First, note that

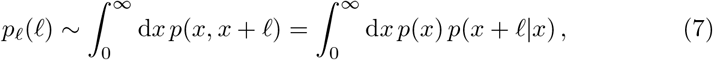

where *p*(*x, x* + *ℓ*) is the joint probability to find one signal at position *x* and another at *x* + *ℓ*, without any signal in between them. *p_ℓ_*(*ℓ*) then averages this quantity over all start positions x. We then express the joint probability of two events as the probability of the first times the probability that the second happens, *given* the first. The result is the probability to find an intersignal distance of *ℓ* anywhere along the (semi-infinite) genome segment.

We also assume an exponentially decreasing amount of label, which implies an exponentially decreasing incorporation rate:

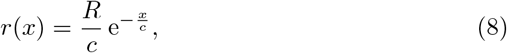

where *c* is the genome distance over which the signal-incorporation rate decreases by a factor e^-1^ ≈ 0.37 and *c/R* is the average distance between two signals at *t* = 0 (i.e., in the absence of depletion). If the fork speed is *v*, then *c/v* is the time it takes for labeled nucleotide concentration to decrease by 37%.

If we assume that the nucleotide concentration correlates with signal probability, then the probability to see a signal at position *x* is also given by

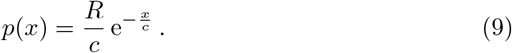

Furthermore, one can check that

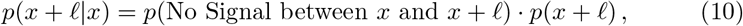

with

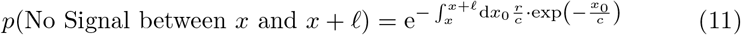

and

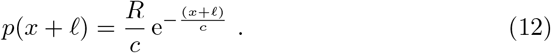

Therefore, one gets

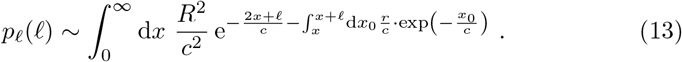

This integral was approximated using Maple (https://www.maplesoft.com) leading to the final result,

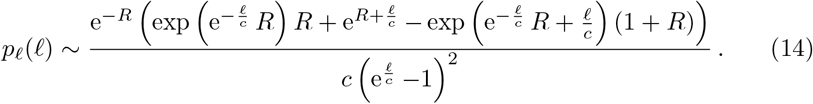

Implicitly, the model above assumes each fiber samples just a single fork whose origin is at *x* = 0. Then *x* is the distance a fork has traveled to the right when the labeled nucleotide (signal) is incorporated. What about more complicated scenarios that a fiber might have? For a single fork moving to the left, the result still holds, as the data reports *unsigned* intersignal distances. Furthermore, numerical results show that the distribution still approximately holds for fibers with multiple forks from neighboring origins.

### 3 Inferring the Position of Initiation

In this section, we describe a method to infer the position at which replication has initiated, given an observed pattern of signals. Assume that one has a segment with signals at positions *x*_1_, *x*_2_,…, *x_n_* (we set *x*_1_ < *x*_2_ < … < *x_n_*). If we assume that the segment was initiated at position *y*, then the probability to observe signals at positions *x*_1_, *x*_2_,…, *x_n_*, is given by

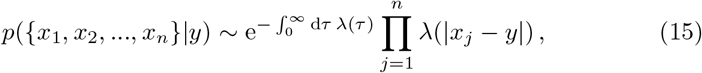

where *λ*(*x*) is the probability to label at a distance *x* from the initiation,

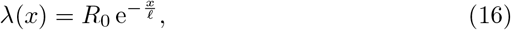

*R*_0_ and *ℓ* being the label and depletion fit parameters. To estimate the position of *y*, we can now do a maximum-likelihood estimation,

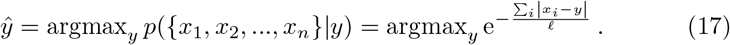

If there is an odd number of signals, then this optimization gives

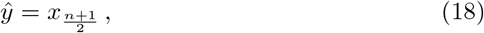

and if there is an even number of signals, the solution is degenerate and can be anything between 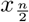 and 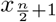. For our calculation, we set

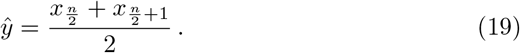

We estimated the uncertainty on this estimator by doing 10^5^ simulations using custom C code (available by request). With the correct fit parameters, this gives us a standard deviation of

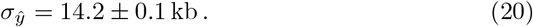

### 4 Estimating the Initiation Event Labeling Efficiency

Using the analysis in Sections 1, 2 and 3, we can estimate the frequency with which an early initiation event will incorporate at least one label and thus be identified by ORM. We begin by noting that the incorporation rate of at position *x* of a replication that started at *t* = 0 is equal to

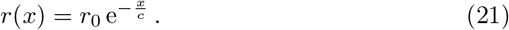

This means that at time *t*, this rate is

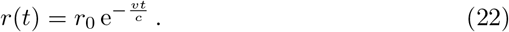

As *r*(*t*) is independent of when the initiation started, one can see that r(*x*) for an initiation that started at time t0 is given by

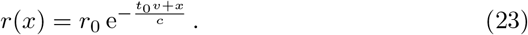

The probability to not get any signals within a distance *x*_0_ from the initiation is then

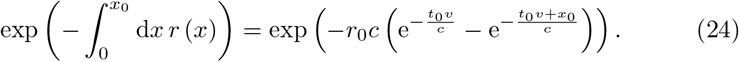

Setting *v*=1.65 kb/min (replication fork rate, from Figure 1C), *x*_0_=15 kb (the nominal resolution of ORM from Eq. 20), *r*_0_=1/3.8 kb (the initial labeling rate, from Figure S2A) and *c*=99 kb (Figure S2b; note that the 75 kb reported there is *c* in base 2, whereas 99 kb used here is *c* in base e) and assuming that the initiations happen uniformly in early S phase, one estimates that the probability to see zero signals within the first 15 kb of an initiation is 9.5%.

### 5 Distribution of Signals within Initiation Zones

Consider an initiation zone of length *L*. We are interested in determining the distribution of initiations inside the initiation zone. Here, we will consider two extreme cases. The first possibility is that the initiation always happens at a single point at the center of the IZ. The second possibility is that the initiation happens with equal probability everywhere along the IZ.

If the initiation always happens at the center of the IZ, then the probability that a signal is incorporated at the center of the IZ, *p_c_*, and the probability that a signal is incorporated at the end of an IZ, *p_e_*, are related via

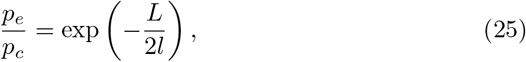

where *l* is the depletion length. On the other hand, if the initiation happens everywhere with equal probability, then the probability to have a signal at an end of the IZ is given by

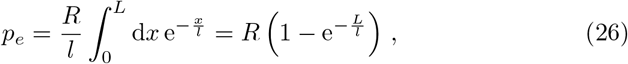

while the probability to have a signal at the center of the IZ is given by

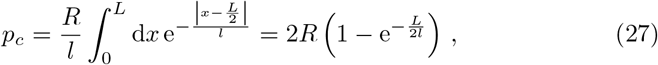

which leads to

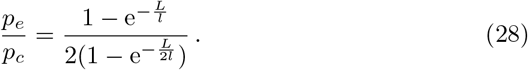

To test whether one of these two models fits the data, we calculate the number of signals within 5 kb of the left end of an IZ (*N_e_*) and the number of signals within 5 kb of the center of the IZ. One then expects

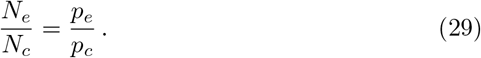

Therefore, we compare *N_e_/N_c_* with Eqs. (25) and (27), where we have determined l from the intersignal distance, *l* = 10^5^.

### 6 Correlation of Labeling in Neighboring Initiation Zones

We consider the correlation function that two neighboring IZ’s, *i* and *i* + 1, are labelled,

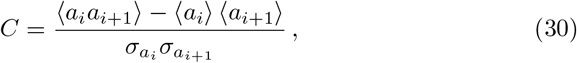

where *α_i_* = 1 if IZ *i* is labeled and 0 if not, and *σ_ai_* is the standard deviation of *a_i_*. As *a_i_* is a binary observable, one can easily check that

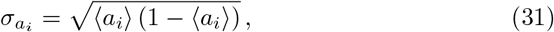

and similarly for *a*_*i*+1_.

Let us assume a model of uncorrelated initiations and, for the moment, that fluctuations in amount of label are irrelevant. In this model, the only correlation between *a_i_* and *a*_*i*+1_ comes from passive replication: if IZ *i* initiates, there is a measurable chance that IZ *i* + 1 gets labelled because it is passively replicated by the fork initiated in IZ *i*. As the amount of label, and therefore the probability to get labelled, decays exponentially with the distance between the IZs, with rate the depletion rate, one would expect that, approximately,

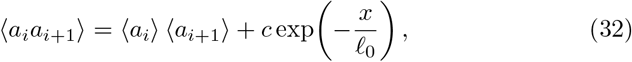

where *c* is an unknown parameter, x is the distance between *i* and *i* + 1 and *ℓ*_0_ is the (1/e) depletion length scale (106 ± 1 kb, Figure S2B).

Let us now include the effect of fluctuations in amount of label between cells. One can write

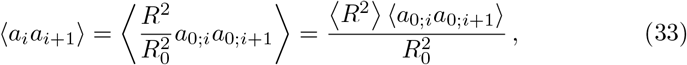

where *R* is the amount of label in the cell, *R*_0_ is the average amount of label in a cell and *a*_0;*i*_ is *a_i_* if the amount of label were to be the average amount of label. If we plug in Eqs. 31, 32 and 33 into Eq. 30, we get

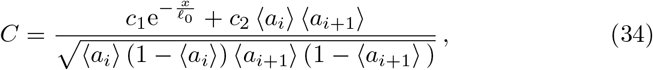

where

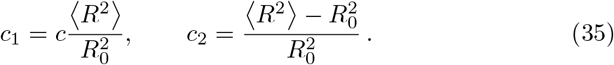

are fit parameters. We leave *c*_1_ as a fit parameter, and from the experimental data, we estimate *c*_2_ = 0.55 ± 0.05 (Figure S1B). Fitting gives

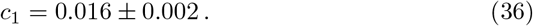

## References

[1] M. Abramowitz, I. Stegun, Eds., Handbook of Mathematical Functions: with Formulas, Graphs, and Mathematical Tables, US Government printing office (1948).

## References

Abdurashidova, G., Deganuto, M., Klima, R., Riva, S., Biamonti, G., Giacca, M., and Falaschi, A. (2000). Start sites of bidirectional DNA synthesis at the human lamin B2 origin. Science 287, 2023–2026.

Anglana, M., Apiou, F., Bensimon, A., and Debatisse, M. (2003). Dynamics of DNA replication in mammalian somatic cells: nucleotide pool modulates origin choice and interorigin spacing. Cell 114, 385–394.

Bechhoefer, J., and Rhind, N. (2012). Replication timing and its emergence from stochastic processes. Trends Genet 28, 374–381.

Bensimon, A., Simon, A., Chiffaudel, A., Croquette, V., Heslot, F., and Bensimon, D. (1994). Alignment and sensitive detection of DNA by a moving interface. Science 265, 2096–2098.

Berezney, R., Dubey, D. D., and Huberman, J. A. (2000). Heterogeneity of eukaryotic replicons, replicon clusters, and replication foci. Chromosoma 108, 471–484.

Bernstein, B. E., Birney, E., Dunham, I., Green, E. D., Gunter, C., and Snyder, M. (2012). An integrated encyclopedia of DNA elements in the human genome. Nature 489, 57–74.

Besnard, E., Babled, A., Lapasset, L., Milhavet, O., Parrinello, H., Dantec, C., Marin, J. M., and Lemaitre, J. M. (2012). Unraveling cell type-specific and reprogrammable human replication origin signatures associated with G-quadruplex consensus motifs. Nat Struct Mol Biol

Birnbaum, A. (1954). Statistical Methods for Poisson Processes and Exponential Populations. Journal of the American Statistical Association 49, 254–266.

Blow, J. J., Gillespie, P. J., Francis, D., and Jackson, D. A. (2001). Replication origins in Xenopus egg extract are 5-15 kilobases apart and are activated in clusters that fire at different times. J Cell Biol 152, 15–25.

Cayrou, C., Ballester, B., Peiffer, I., Fenouil, R., Coulombe, P., Andrau, J. C., van Helden, J., and Mechali, M. (2015). The chromatin environment shapes DNA replication origin organization and defines origin classes. Genome Res 25, 1873–1885.

Cayrou, C., Coulombe, P., Vigneron, A., Stanojcic, S., Ganier, O., Peiffer, I., Rivals, E., Puy, A., Laurent-Chabalier, S., Desprat, R., and Mechali, M. (2011). Genome-scale analysis of metazoan replication origins reveals their organization in specific but flexible sites defined by conserved features. Genome Res 21, 1438–1449.

Chagin, V. O., Casas-Delucchi, C. S., Reinhart, M., Schermelleh, L., Markaki, Y., Maiser, A., Bolius, J. J., Bensimon, A., Fillies, M., Domaing, P., Rozanov, Y. M., Leonhardt, H., and Cardoso, M. C. (2016). 4D Visualization of replication foci in mammalian cells corresponding to individual replicons. Nat Commun 7, 11231.

Chen, C. L., Duquenne, L., Audit, B., Guilbaud, G., Rappailles, A., Baker, A., Huvet, M., d’Aubenton-Carafa, Y., Hyrien, O., Arneodo, A., and Thermes, C. (2011). Replication-associated mutational asymmetry in the human genome. Mol Biol Evol 28, 2327–2337.

Chen, C. L., Rappailles, A., Duquenne, L., Huvet, M., Guilbaud, G., Farinelli, L., Audit, B., d’Aubenton-Carafa, Y., Arneodo, A., Hyrien, O., and Thermes, C. (2010). Impact of replication timing on non-CpG and CpG substitution rates in mammalian genomes. Genome Res 20, 447–457.

Conti, C., Saccà, B., Herrick, J., Lalou, C., Pommier, Y., and Bensimon, A. (2007). Replication fork velocities at adjacent replication origins are coordinately modified during DNA replication in human cells. Mol Biol Cell 18, 3059–3067.

Czajkowsky, D. M., Liu, J., Hamlin, J. L., and Shao, Z. (2008). DNA Combing Reveals Intrinsic Temporal Disorder in the Replication of Yeast Chromosome VI. J Mol Biol 375, 12–19.

De Carli, F., Menezes, N., Berrabah, W., Barbe, V., Genovesio, A., and Hyrien, O. (2018). High-Throughput Optical Mapping of Replicating DNA. Small Methods 0, 1800146.

de Moura, A. P., Retkute, R., Hawkins, M., and Nieduszynski, C. A. (2010). Mathematical modelling of whole chromosome replication. Nucleic Acids Res 38, 5623–5633.

Dellino, G. I., Cittaro, D., Piccioni, R., Luzi, L., Banfi, S., Segalla, S., Cesaroni, M., Mendoza-Maldonado, R., Giacca, M., and Pelicci, P. G. (2013). Genome-wide mapping of human DNA-replication origins: levels of transcription at ORC1 sites regulate origin selection and replication timing. Genome Res 23, 1–11.

Demczuk, A., Gauthier, M. G., Veras, I., Kosiyatrakul, S., Schildkraut, C. L., Busslinger, M., Bechhoefer, J., and Norio, P. (2012). Regulation of DNA Replication within the Immunoglobulin Heavy-Chain Locus During B Cell Commitment. PLoS Biol 10, e1001360.

Dijkwel, P. A., Wang, S., and Hamlin, J. L. (2002). Initiation sites are distributed at frequent intervals in the Chinese hamster dihydrofolate reductase origin of replication but are used with very different efficiencies. Mol Cell Biol 22, 3053–3065.

Dileep, V., and Gilbert, D. M. (2018). Single-cell replication profiling to measure stochastic variation in mammalian replication timing. Nat Commun 9, 427.

Eaton, M. L., Galani, K., Kang, S., Bell, S. P., and Macalpine, D. M. (2010). Conserved nucleosome positioning defines replication origins. Genes Dev 24, 748–753.

Edwards, M. C., Tutter, A. V., Cvetic, C., Gilbert, C. H., Prokhorova, T. A., and Walter, J. C. (2002). MCM2-7 complexes bind chromatin in a distributed pattern surrounding the origin recognition complex in Xenopus egg extracts. J Biol Chem 277, 33049–33057.

Ernst, J., and Kellis, M. (2012). ChromHMM: automating chromatin-state discovery and characterization. Nat Methods 9, 215–216.

Farkash-Amar, S., Lipson, D., Polten, A., Goren, A., Helmstetter, C., Yakhini, Z., and Simon, I. (2008). Global organization of replication time zones of the mouse genome. Genome Res 18, 1562–1570.

Feijoo, C., Hall-Jackson, C., Wu, R., Jenkins, D., Leitch, J., Gilbert, D. M., and Smythe, C. (2001). Activation of mammalian Chk1 during DNA replication arrest: a role for Chk1 in the intra-S phase checkpoint monitoring replication origin firing. J Cell Biol 154, 913–923.

Foulk, M. S., Urban, J. M., Casella, C., and Gerbi, S. A. (2015). Characterizing and controlling intrinsic biases of lambda exonuclease in nascent strand sequencing reveals phasing between nucleosomes and G-quadruplex motifs around a subset of human replication origins. Genome Res 25, 725–735.

Fragkos, M., Ganier, O., Coulombe, P., and Méchali, M. (2015). DNA replication origin activation in space and time. Nat Rev Mol Cell Biol 16, 360–374.

Ganier, O., Prorok, P., Akerman, I., and Méchali, M. (2019). Metazoan DNA replication origins. Curr Opin Cell Biol 58, 134–141.

Georgieva, D., Liu, Q., Wang, K., and Egli, D. (2019). Detection of Base Analogs Incorporated During DNA Replication by Nanopore Sequencing. bioRxiv 549220.

Gindin, Y., Meltzer, P. S., and Bilke, S. (2014a). Replicon: a software to accurately predict DNA replication timing in metazoan cells. Front Genet 5, 378.

Gindin, Y., Valenzuela, M. S., Aladjem, M. I., Meltzer, P. S., and Bilke, S. (2014b). A chromatin structure-based model accurately predicts DNA replication timing in human cells. Mol Syst Biol 10, 722.

Goldar, A., Marsolier-Kergoat, M. C., and Hyrien, O. (2009). Universal temporal profile of replication origin activation in eukaryotes. PLoS One 4, e5899.

Gros, J., Kumar, C., Lynch, G., Yadav, T., Whitehouse, I., and Remus, D. (2015). Post-licensing Specification of Eukaryotic Replication Origins by Facilitated Mcm2-7 Sliding along DNA. Mol Cell 60, 797–807.

Guilbaud, G., Rappailles, A., Baker, A., Chen, C. L., Arneodo, A., Goldar, A., d’Aubenton-Carafa, Y., Thermes, C., Audit, B., and Hyrien, O. (2011). Evidence for Sequential and Increasing Activation of Replication Origins along Replication Timing Gradients in the Human Genome. PLoS Comput Biol 7, e1002322.

Hamlin, J. L., Mesner, L. D., Lar, O., Torres, R., Chodaparambil, S. V., and Wang, L. (2008). A revisionist replicon model for higher eukaryotic genomes. J Cell Biochem 105, 321–329.

Hansen, R. S., Thomas, S., Sandstrom, R., Canfield, T. K., Thurman, R. E., Weaver, M., Dorschner, M. O., Gartler, S. M., and Stamatoyannopoulos, J. A. (2010). Sequencing newly replicated DNA reveals widespread plasticity in human replication timing. Proc Natl Acad Sci U S A 107, 139–144.

Hennion, M., Arbona, J. M., Lacroix, L., Cruaud, C., Theulot, B., Tallec, B. L., Proux, F., Wu, X., Novikova, E., Engelen, S., Lemainque, A., Audit, B., and Hyrien, O. (2020). FORK-seq: replication landscape of the Saccharomyces cerevisiae genome by nanopore sequencing. Genome Biol 21, 125.

Herrick, J., and Bensimon, A. (1999). Single molecule analysis of DNA replication. Biochimie 81, 859–871.

Hiratani, I., Ryba, T., Itoh, M., Yokochi, T., Schwaiger, M., Chang, C. W., Lyou, Y., Townes, T. M., Schubeler, D., and Gilbert, D. M. (2008). Global reorganization of replication domains during embryonic stem cell differentiation. PLoS Biol 6, e245.

Huberman, J. A., and Riggs, A. D. (1968). On the mechanism of DNA replication in mammalian chromosomes. J Mol Biol 32, 327–341.

Huberman, J. A., and Tsai, A. (1973). Direction of DNA replication in mammalian cells. J Mol Biol 75, 5–12.

Hyrien, O. (2015). Peaks cloaked in the mist: the landscape of mammalian replication origins. J Cell Biol 208, 147–160.

Jackson, D. A., and Pombo, A. (1998). Replicon clusters are stable units of chromosome structure: evidence that nuclear organization contributes to the efficient activation and propagation of S phase in human cells. J Cell Biol 140, 1285–1295.

Kaykov, A., and Nurse, P. (2015). The spatial and temporal organization of origin firing during the S-phase of fission yeast. Genome Res 25, 391–401.

Kaykov, A., Taillefumier, T., Bensimon, A., and Nurse, P. (2016). Molecular Combing of Single DNA Molecules on the 10 Megabase Scale. Sci Rep 6, 19636.

Keller, C., Ladenburger, E. M., Kremer, M., and Knippers, R. (2002). The origin recognition complex marks a replication origin in the human TOP1 gene promoter. J Biol Chem 277, 31430–31440.

Lam, E. T., Hastie, A., Lin, C., Ehrlich, D., Das, S. K., Austin, M. D., Deshpande, P., Cao, H., Nagarajan, N., Xiao, M., and Kwok, P. Y. (2012). Genome mapping on nanochannel arrays for structural variation analysis and sequence assembly. Nat Biotechnol 30, 771–776.

Langley, A. R., Gräf, S., Smith, J. C., and Krude, T. (2016). Genome-wide identification and characterisation of human DNA replication origins by initiation site sequencing (ini-seq). Nucleic Acids Res 44, 10230–10247.

Lebofsky, R., Heilig, R., Sonnleitner, M., Weissenbach, J., and Bensimon, A. (2006). DNA Replication Origin Interference Increases the Spacing between Initiation Events in Human Cells. Mol Biol Cell 17, 5337–5345.

Löb, D., Lengert, N., Chagin, V. O., Reinhart, M., Casas-Delucchi, C. S., Cardoso, M. C., and Drossel, B. (2016). 3D replicon distributions arise from stochastic initiation and domino-like DNA replication progression. Nat Commun 7, 11207.

Long, H., Zhang, L., Lv, M., Wen, Z., Zhang, W., Chen, X., Zhang, P., Li, T., Chang, L., Jin, C., Wu, G., Wang, X., Yang, F., Pei, J., Chen, P., Margueron, R., Deng, H., Zhu, M., and Li, G. (2020). H2A.Z facilitates licensing and activation of early replication origins. Nature 577, 576–581.

Lubelsky, Y., Sasaki, T., Kuipers, M. A., Lucas, I., Le Beau, M. M., Carignon, S., Debatisse, M., Prinz, J. A., Dennis, J. H., and Gilbert, D. M. (2011). Pre-replication complex proteins assemble at regions of low nucleosome occupancy within the Chinese hamster dihydrofolate reductase initiation zone. Nucleic Acids Res 39, 3141–3155.

Macheret, M., and Halazonetis, T. D. (2018). Intragenic origins due to short G1 phases underlie oncogene-induced DNA replication stress. Nature 555, 112–116.

Marchal, C., Sima, J., and Gilbert, D. M. (2019). Control of DNA replication timing in the 3D genome. Nat Rev Mol Cell Biol 20, 721–737.

Marheineke, K., and Hyrien, O. (2004). Control of replication origin density and firing time in Xenopus egg extracts: role of a caffeine-sensitive, ATR-dependent checkpoint. J Biol Chem 279, 28071–28081.

Masai, H., Matsumoto, S., You, Z., Yoshizawa-Sugata, N., and Oda, M. (2010). Eukaryotic chromosome DNA replication: where, when, and how? Annu Rev Biochem 79, 89–130.

McGuffee, S. R., Smith, D. J., and Whitehouse, I. (2013). Quantitative, genome-wide analysis of eukaryotic replication initiation and termination. Mol Cell 50, 123–135.

Mesner, L. D., Li, X., Dijkwel, P. A., and Hamlin, J. L. (2003). The dihydrofolate reductase origin of replication does not contain any nonredundant genetic elements required for origin activity. Mol Cell Biol 23, 804–814.

Mesner, L. D., Valsakumar, V., Cieslik, M., Pickin, R., Hamlin, J. L., and Bekiranov, S. (2013). Bubble-seq analysis of the human genome reveals distinct chromatin-mediated mechanisms for regulating early- and late-firing origins. Genome Res 23, 1774–1788.

Miotto, B., Ji, Z., and Struhl, K. (2016). Selectivity of ORC binding sites and the relation to replication timing, fragile sites, and deletions in cancers. Proc Natl Acad Sci U S A 113, E4810–9.

Mitter, M., Gasser, C., Takacs, Z., Langer, C. C. H., Tang, W., Jessberger, G., Beales, C. T., Neuner, E., Ameres, S. L., Peters, J. M., Goloborodko, A., Micura, R., and Gerlich, D. W. (2020). Conformation of sister chromatids in the replicated human genome. Nature 586, 139–144.

Müller, C. A., Boemo, M. A., Spingardi, P., Kessler, B. M., Kriaucionis, S., Simpson, J. T., and Nieduszynski, C. A. (2019). Capturing the dynamics of genome replication on individual ultra-long nanopore sequence reads. Nat Methods 442814.

Neef, A. B., and Luedtke, N. W. (2014). An azide-modified nucleoside for metabolic labeling of DNA. Chembiochem 15, 789–793.

Norio, P., and Schildkraut, C. L. (2001). Visualization of DNA replication on individual Epstein-Barr virus episomes. Science 294, 2361–2364.

O’Keefe, R. T., Henderson, S. C., and Spector, D. L. (1992). Dynamic organization of DNA replication in mammalian cell nuclei: spatially and temporally defined replication of chromosome-specific alphasatellite DNA sequences. J Cell Biol 116, 1095–1110.

Panning, M. M., and Gilbert, D. M. (2005). Spatio-temporal organization of DNA replication in murine embryonic stem, primary, and immortalized cells. J Cell Biochem 95, 74–82.

Patel, P. K., Arcangioli, B., Baker, S. P., Bensimon, A., and Rhind, N. (2006). DNA replication origins fire stochastically in fission yeast. Mol Biol Cell 17, 308–316.

Petryk, N., Kahli, M., d’Aubenton-Carafa, Y., Jaszczyszyn, Y., Shen, Y., Silvain, M., Thermes, C., Chen, C. L., and Hyrien, O. (2016). Replication landscape of the human genome. Nat Commun 7, 10208.

Picard, F., Cadoret, J. C., Audit, B., Arneodo, A., Alberti, A., Battail, C., Duret, L., and Prioleau, M. N. (2014). The spatiotemporal program of DNA replication is associated with specific combinations of chromatin marks in human cells. PLoS Genet 10, e1004282.

Pope, B. D., Ryba, T., Dileep, V., Yue, F., Wu, W., Denas, O., Vera, D. L., Wang, Y., Hansen, R. S., Canfield, T. K., Thurman, R. E., Cheng, Y., Gulsoy, G., Dennis, J. H., Snyder, M. P., Stamatoyannopoulos, J. A., Taylor, J., Hardison, R. C., Kahveci, T., Ren, B., and Gilbert, D. M. (2014). Topologically associating domains are stable units of replication-timing regulation. Nature 515, 402–405.

Pourkarimi, E., Bellush, J. M., and Whitehouse, I. (2016). Spatiotemporal coupling and decoupling of gene transcription with DNA replication origins during embryogenesis in C. elegans. Elife 5,

Powell, S. K., MacAlpine, H. K., Prinz, J. A., Li, Y., Belsky, J. A., and MacAlpine, D. M. (2015). Dynamic loading and redistribution of the Mcm2-7 helicase complex through the cell cycle. EMBO J 34, 531–543.

Prorok, P., Artufel, M., Aze, A., Coulombe, P., Peiffer, I., Lacroix, L., Guédin, A., Mergny, J. L., Damaschke, J., Schepers, A., Cayrou, C., Teulade-Fichou, M. P., Ballester, B., and Méchali, M. (2019). Involvement of G-quadruplex regions in mammalian replication origin activity. Nat Commun 10, 3274.

Puig Lombardi, E., Holmes, A., Verga, D., Teulade-Fichou, M. P., Nicolas, A., and Londoño-Vallejo, A. (2019). Thermodynamically stable and genetically unstable G-quadruplexes are depleted in genomes across species. Nucleic Acids Res 47, 6098–6113.

Ramírez, F., Ryan, D. P., Grüning, B., Bhardwaj, V., Kilpert, F., Richter, A. S., Heyne, S., Dündar, F., and Manke, T. (2016). deepTools2: a next generation web server for deep-sequencing data analysis. Nucleic Acids Res 44, W160–5.

Rhind, N., and Gilbert, D. M. (2013). DNA replication timing. Cold Spring Harb Perspect Biol 5, a010132.

Rhind, N., Yang, S. C.-H., and Bechhoefer, J. (2010). Reconciling stochastic origin firing with defined replication timing. Chromosome Res 18, 35–43.

Rieder, U., and Luedtke, N. W. (2014). Alkene-Tetrazine Ligation for Imaging Cellular DNA. Angew Chem Int Ed Engl 53, 9168–9172.

Robin, X., Turck, N., Hainard, A., Tiberti, N., Lisacek, F., Sanchez, J. C., and Müller, M. (2011). pROC: an open-source package for R and S+ to analyze and compare ROC curves. BMC Bioinformatics 12, 77.

Robinson, J. T., Thorvaldsdóttir, H., Winckler, W., Guttman, M., Lander, E. S., Getz, G., and Mesirov, J. P. (2011). Integrative genomics viewer. Nat Biotechnol 29, 24–26.

Tao, L., Dong, Z., Leffak, M., Zannis-Hadjopoulos, M., and Price, G. (2000). Major DNA replication initiation sites in the c-myc locus in human cells. J Cell Biochem 78, 442–457.

Taylor, J. H. (1977). Increase in DNA replication sites in cells held at the beginning of S phase. Chromosoma 62, 291–300.

Técher, H., Koundrioukoff, S., Azar, D., Wilhelm, T., Carignon, S., Brison, O., Debatisse, M., and Le Tallec, B. (2013). Replication dynamics: biases and robustness of DNA fiber analysis. J Mol Biol 425, 4845–4855.

Theis, J. F., and Newlon, C. S. (1997). The ARS309 chromosomal replicator of Saccharomyces cerevisiae depends on an exceptional ARS consensus sequence. Proc Natl Acad Sci U S A 94, 10786–10791.

Valton, A. L., Hassan-Zadeh, V., Lema, I., Boggetto, N., Alberti, P., Saintome, C., Riou, J. F., and Prioleau, M. N. (2014). G4 motifs affect origin positioning and efficiency in two vertebrate replicators. EMBO J 33, 732–746.

Vashee, S., Cvetic, C., Lu, W., Simancek, P., Kelly, T. J., and Walter, J. C. (2003). Sequence-independent DNA binding and replication initiation by the human origin recognition complex. Genes Dev 17, 1894–1908.

Wilson, K. A., Elefanty, A. G., Stanley, E. G., and Gilbert, D. M. (2016). Spatio-temporal re-organization of replication foci accompanies replication domain consolidation during human pluripotent stem cell lineage specification. Cell Cycle 15, 2464–2475.

Xu, J., Yanagisawa, Y., Tsankov, A. M., Hart, C., Aoki, K., Kommajosyula, N., Steinmann, K. E., Bochicchio, J., Russ, C., Regev, A., Rando, O. J., Nusbaum, C., Niki, H., Milos, P., Weng, Z., and Rhind, N. (2012). Genome-wide identification and characterization of replication origins by deep sequencing. Genome Biol 13, R27.

Yang, S. C.-H., and Bechhoefer, J. (2008). How Xenopus laevis embryos replicate reliably: Investigating the random-completion problem. Phys Rev E Stat Nonlin Soft Matter Phys 78, 041917.

Yang, S. C.-H., Rhind, N., and Bechhoefer, J. (2010). Modeling genome-wide replication kinetics reveals a mechanism for regulation of replication timing. Mol Syst Biol 6, 404.

Zhao, P. A., Sasaki, T., and Gilbert, D. M. (2020). High-resolution Repli-Seq defines the temporal choreography of initiation, elongation and termination of replication in mammalian cells. Genome Biol 21, 76.

